# Plant growth forms shaping biodiversity patterns can guide conservation planning on the Qinghai-Tibet Plateau

**DOI:** 10.64898/2026.05.31.729149

**Authors:** Feng Liu, Zhiyuan Liang, Xiao Xu, Jingge Kuang, Jin Ye, Filing Yang, Jie Li, Yupeng Geng, Bo Li, Jinming Hu

## Abstract

Plant growth forms effectively reflect profound evolutionary adaptations that underpin ecosystem functioning; however, how their biogeographic disparities influence conservation prioritization remains poorly understood. We here integrate a comprehensive phylogeny with high-resolution distribution data for 14,468 vascular plant species across the Qinghai-Tibet Plateau, showing that growth-form identity decouples the spatial patterns of taxonomic and phylogenetic diversity, with distinct forms exhibiting marked differences in geographic range sizes and elevational optima (2,200–3,300 m). Although herbs dominate plant assemblages across spatial scales, their proportional abundance decreases along a northwest-to-southeast aridity gradient, yielding to structurally complex, non-herbaceous forms in humid, species-rich regions. Consequently, priority conservation areas derived for individual growth forms exhibit low spatial similarity. Despite this divergence, we identify trees and climbers as highly effective umbrella growth forms: prioritizing these functional groups captures ≥75% of the multidimensional diversity of all other growth forms within a minimal spatial footprint. Our findings demonstrate that accounting for growth-form-based functional identity is essential for maximizing conservation efficacy, providing a scalable framework for biodiversity hotspots worldwide.

## 1. Introduction

Effective conservation planning relies on a robust understanding of biodiversity patterns to prioritize limited resources strategically [1–4]. To navigate the complexity of biodiversity, conservationists often adopt surrogacy—using well-studied taxa to represent broader diversity [5–6]. However, the efficacy of this approach is frequently undermined by spatial mismatches among taxonomic groups [4,7]. Functional biogeography resolves these discrepancies by revealing their mechanistic basis: plant growth forms (e.g., trees, shrubs, and herbs) are not merely morphological categories, but represent fundamental, distinct functional strategies [8]. As convergent evolutionary solutions to physiological constraints, these growth forms interact with the environment in divergent ways [8–9]. Consequently, they serve as powerful, scalable surrogates for conservation prioritization, capturing the functional scaffolding of ecosystems beyond simple species counts [2,8–10]. Relying solely on broad taxonomic surrogates treats plant diversity as a monolith, systematically masking the distinct spatial distributions inherent to these vital functional groups [9,11–12].

This distributional specificity is rooted in fundamental ecophysiological divergence and evolutionary conservatism [9–10]. For instance, constrained by tropical origins and hydraulic vulnerability, tree lineages often remain restricted to stable, warmer lowlands [5,9], whereas climbers and epiphytes exploit the complex structural niches of tropical montane forests [13–14]. Conversely, shrubs and herbs dominate more stressful environments: shrubs employ sclerophylly to inhabit climatically variable and arid zones [10], while herbs utilize flexible life-history strategies to colonize high-altitude or xeric niches [15]. These ecological disparities are entrenched by phylogenetic niche conservatism, which restricts environmental adaptation and clusters specific functional strategies (e.g., epiphytism in Orchidaceae) within distinct lineages [16–17]. Consequently, even in sympatry, different growth forms may constitute compositionally distinct assemblages driven by contrasting ecological processes [10].

Despite these fundamental ecological distinctions, the extent to which relying on broad taxonomic surrogates distorts conservation prioritization remains largely unquantified. Current paradigms typically depend on aggregate diversity metrics, which obscure the unique adaptive strategies and spatial patterns of distinct growth forms [9,11]. Consequently, integrating functional identity into conservation frameworks is imperative to reassess the efficacy of existing surrogacy approaches and ensure representation of the full spectrum of plant biodiversity.

The Qinghai–Tibet Plateau (QTP) serves as an ideal natural laboratory and a globally significant model system to address this challenge. Spanning ∼2.74 million km² with a mean elevation of ∼4400 m [18] (Fig. 1a), the QTP compresses immense climatic gradients into a complex topography, fostering exceptional endemism across four global biodiversity hotspots [18–19] (Fig. 1b). By compressing global biomes within a single region, the plateau amplifies the functional sorting mechanisms that may remain cryptic in less heterogeneous landscapes [20] (Fig. 1c). As the Third Pole and a sentinel for climate change, this region exemplifies the conservation dilemmas faced by montane ecosystems worldwide. Consequently, resolving functional mismatches here provides a scalable blueprint for safeguarding biodiversity in topographically complex environments globally. In this study we leverage this system to quantify the spatial incongruence among plant growth forms, revealing how distinct functional strategies compromise the effectiveness of traditional surrogacy and necessitate a targeted expansion of conservation priorities.

**Fig. 1.**
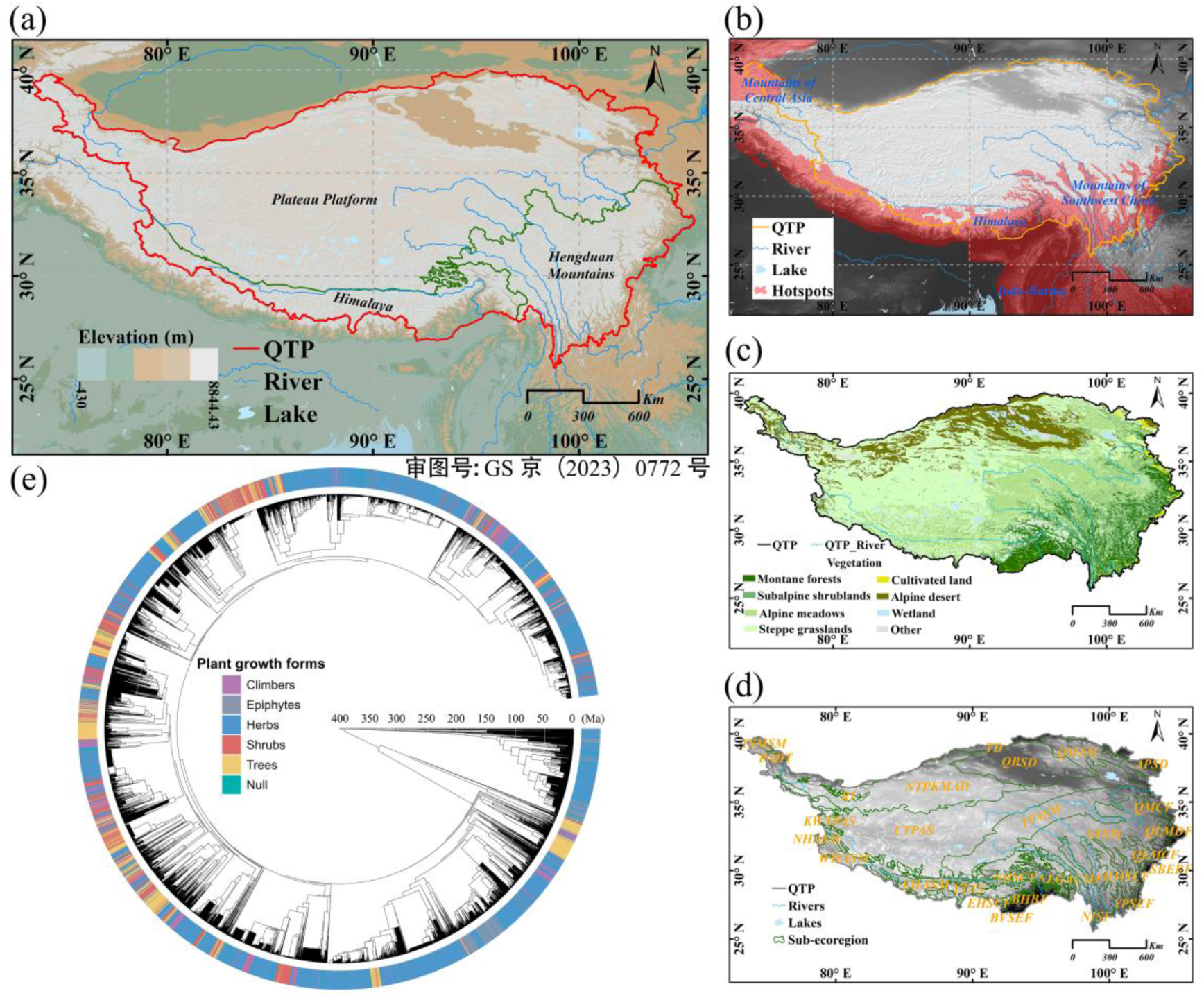
Geographic context and phylogenetic framework of vascular plants on the Qinghai–Tibet Plateau (QTP). (a) Topographic map of the QTP, highlighting its major physiographic units: the Plateau Platform, Hengduan Mountains, and Himalayas. (b) Location of the QTP relative to (and overlapping with) four global biodiversity hotspots (red): the Mountains of Central Asia, the Himalayas, Indo-Burma, and the Mountains of Southwest China. (c) Spatial distribution of dominant vegetation types, illustrating the pronounced ecological gradient from montane forests to alpine deserts. (d) Delineation of sub–ecoregions used for spatial analysis. (e) Phylogenetic tree of the 17,844 vascular plant species recorded across the QTP (serving as the backbone for the 14,468 species analyzed). Terminal branches are colored by growth form: climbers (purple), epiphytes (grey), herbs (blue), shrubs (red), trees (yellow), and species with unresolved growth forms (green; labeled “Null”).

Here, we synthesized an extensive dataset of 14,468 vascular plant species—integrating verified occurrences, growth-form assignments, and phylogenetic data—to construct high-resolution (1 km × 1 km) diversity maps across the QTP using ensemble ecological niche modeling. Through this integrated framework, we address the following questions: (1) To what extent do the spatial patterns of taxonomic and phylogenetic diversity differ across plant growth forms? (2) How does the proportional composition of plant growth forms vary geographically across the QTP? and (3) Which specific plant growth forms are the most effective surrogates for representing others in spatial conservation prioritization? To answer these questions and evaluate the conservation implications of growth form–specific distributions we assessed the spatial congruence of diversity metrics between overall vascular plants (all plants) and individual growth forms, mapped the geographical variation in their composition, and quantified the spatial similarity of priority conservation areas (PCAs) to determine the cross-taxon efficacy of specific growth forms as surrogates.

## 2. Materials and methods

### 2.1 Plant Growth Form and Occurrence Records

We categorized the vascular flora of the Qinghai-Tibet Plateau (QTP) into five functional growth forms: climbers, epiphytes, herbs, shrubs, and trees, following established definitions [10]. Baseline taxonomic classifications were derived primarily from the Flora Reipublicae Popularis Sinicae and the Flora of China. To ensure accuracy and comprehensive coverage, we cross-referenced these initial assignments against specialized datasets, including the Growth-Form Database for Chinese Vascular Plants [21], WCVP 2.0 [22], and the GIFT v3.0 database [23]. In cases of discrepancy, we prioritized regional specialized databases [21], followed by consensus among global repositories. For climbers and epiphytes, we integrated supplementary records from taxon-specific checklists [24–26]. The final curated inventory encompassed 17,283 species (representing approximately 97% of the known QTP flora) across 2,424 genera and 271 families (Supplementary Method 1 and Dataset S1).

To prevent ecological niche truncation in our species distribution models [27], we compiled an occurrence dataset that extended beyond the QTP into adjacent ecoregions (Fig. 1d). We aggregated records from multiple repositories: the Global Biodiversity Information Facility (GBIF; 660,220 records), iDigBio (82,867 records), the Botanical Information and Ecology Network (BIEN; 97,206 observations), the China Virtual Herbarium (28,948 records), the Herbarium of the Institute of Botany, Chinese Academy of Sciences (15,217 specimens), and primary field survey data from the Second Qinghai-Tibet Plateau Expedition. Following a multi-step taxonomic and spatial data cleaning pipeline (Supplementary Method 2), we retained a high-quality dataset of 325,216 georeferenced records representing 14,468 species. This final dataset comprised 8,936 herbs, 2,088 shrubs, 1,776 trees, 1,061 climbers, and 607 epiphytes (Table S1).

### 2.2 Environmental Data

We sourced contemporary climate data (1981–2010) from Climatologies at High Resolution for the Earth’s Land Surface Areas (CHELSA v2.1), extracting 19 bioclimatic variables and growing degree days (GDD). Topographic data were derived from EarthEnv-DEM90 at 30-m resolution. To quantify habitat heterogeneity, we calculated the elevation range and its standard deviation within 1-km cells, alongside the terrain roughness index from the ENVIREM database [28]. Soil properties for the upper 20 cm—specifically total nitrogen, clay content, and total exchangeable bases—were obtained from the Harmonized World Soil Database v2.0 [29], representing key edaphic predictors of plant distributions [6]. All environmental layers were resampled to a consistent 30-arcsecond (∼1 km) resolution.

### 2.3 Mapping species range

To generate standardized, high-resolution range maps, we employed an ensemble species distribution modeling (SDM) framework [4,6] using the biomod2 R package (v.4.2.2). For the 9,637 species with ≥ 5 unique occurrence records, we applied five algorithms: classification tree analysis (CTA), generalized additive models (GAM), generalized boosted regression models (GBM), random forest (RF), and maximum entropy (MaxEnt).

To minimize multicollinearity, we selected species-specific subsets of environmental predictors by excluding variables with a pairwise Pearson correlation > 0.7 or a variance inflation factor (VIF) > 5 [30]. Models were calibrated against 10,000 randomly sampled background points [6,31]. Performance was evaluated using 5-fold cross-validation (80% training, 20% testing) repeated five times per algorithm, yielding 25 initial models per species. We assessed model accuracy using three complementary metrics: Area Under the Curve (AUC), True Skill Statistic (TSS), and Boyce Index [32]. Only models meeting stringent thresholds (AUC ≥ 0.7, TSS ≥ 0.5, and Boyce Index ≥ 0.4) were retained for ensemble forecasting (Table S2). For each species, a continuous habitat suitability map was generated by averaging projections from all qualified models using the EMca algorithm. These continuous maps were then converted to binary presence-absence projections using the MaxTSS threshold. To mitigate potential overprediction [33], the binary maps were spatially constrained to a 200-km buffer surrounding a minimum convex polygon (MCP) defined by the species’ occurrences.

For the remaining 5,812 species—those with fewer than five occurrences or whose models failed to meet performance thresholds—we estimated ranges using a 10-km circular buffer around each occurrence point, following standard protocols for data-poor taxa in macroecological analyses [12]. All final range maps were standardized to a 1-km grid. The resulting dataset comprises 8,656 SDM-derived and 5,812 buffer-derived range maps, collectively capturing 14,468 species (81.1% of the known QTP vascular flora; Table S3).

### 2.4 Diversity metrics

To quantify spatial biodiversity patterns, we calculated four complementary metrics from the high-resolution range maps. Total vascular plant species richness (SR) was derived by overlaying species distribution ranges and summing all co-occurring species within each 1-km grid cell:

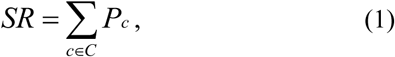

Here, 𝑃_𝑐_ indicates the presence of species 𝑐 in a given region (e.g., a 1-km² grid cell), and 𝐶 denotes the set of all species co-occurring within that region (i.e., the local plant assemblage). Growth form richness was quantified similarly as the cumulative count of species per specific functional group. Additionally, to assess the range size of each group, we calculated the total occupied area (in km²) by summing the areas of all grid cells where the corresponding richness exceeded zero.

Weighted endemism (WE), a widely employed metric for identifying centers of endemism, quantifies species’ range restrictions through inverse weighting by their distributional extent [34]:

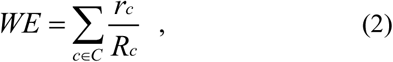

*r_c_* denotes the localised distribution range of species c (operationally defined as 1-km² grid cell), *R_c_* represents the total geographic range of species c, and C corresponds to the set of species within the given grid cell.

To quantify evolutionary metrics, we reconstructed a comprehensive phylogeny using the plant mega-phylogeny GBOTB.extended.WP.tre [35] as the backbone. For species absent from this backbone, we employed the U.PhyloMaker R package [36] to insert them as polytomies at their corresponding genus or family nodes using ‘Scenario S3’. We then pruned this complete phylogeny (Fig. 1e) to retain only the 14,468 species with valid distribution data, forming the core phylogeny for assemblage-level analyses (*Wightia speciosissima* was excluded due to taxonomic uncertainty). To investigate patterns across functional groups, we generated five growth-form-specific sub-phylogenies by pruning the core tree: herbs (8,936 species; 4,373 internal nodes), shrubs (2,088 species; 1,159 internal nodes), trees (1,775 species; 1,205 internal nodes), climbers (1,061 species; 760 internal nodes), and epiphytes (607 species; 312 internal nodes).

Phylogenetic diversity (PD) was quantified as Faith’s PD [37], representing the total branch length of the minimal spanning subtree that includes all species in an assemblage:

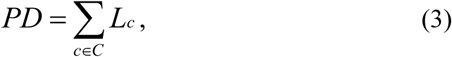

where C represents the set of branches within a given region, and *L_c_* indicates the branch length of *c*.

Phylogenetic endemism (PE) measures the spatial restriction of evolutionary history by summing branch lengths inversely weighted by their spatial occupancy:

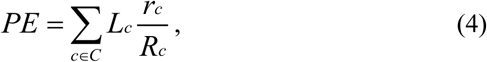

Within Equation (4), *C* denotes the set of branches in a given region. *L_c_* the branch length of *c*. *r_c_* is localised distribution range (measured as 1 km² grid cell), and *R_c_* signifies the total geographic range size of the branch.

Finally, we calculated all biodiversity metrics (SR, WE, PD, and PE) for all plants and each growth form separately using the phyloregion R package [38]. To characterize large-scale biogeographic patterns, we modeled the relationships between each diversity metric and key environmental variables (latitude and elevation) using ordinary least squares (OLS) regression with cubic polynomials to account for potential non-linear responses. Furthermore, spatial congruence among diversity patterns—both overall and across discrete growth forms—was assessed using pairwise Spearman’s rank correlation coefficients across all grid cells.

### 2.5 Compositional structure of plant growth forms

We quantified growth-form composition across both taxonomic and phylogenetic dimensions. Taxonomically, this was expressed as the relative species richness of each growth form (i.e., the ratio of growth-form-specific richness to total vascular plant richness). Phylogenetically, we measured the relative representation of each growth form as the ratio of its specific phylogenetic diversity (PD) to the total PD of the entire assemblage. We evaluated these metrics hierarchically across three spatial scales: (1) regional (the entire QTP), utilizing the 17,283 species with assigned growth forms from the initial checklist; (2) subregional (major physiographic units and sub-ecoregions [39]; Fig. 1d), utilizing the aggregated range maps of the 14,468 mapped species; and (3) local (within each 1-km² grid cell), based on the co-occurring species assemblages derived from our high-resolution range maps.

### 2.6 Spatial conservation prioritization across growth forms

To identify taxonomic and phylogenetic conservation priorities, we employed the spatial prioritization framework Zonation 5.0 [40]. Our analysis integrated two distinct biodiversity features: (1) species-level range maps for all 14,468 vascular plants, and (2) the spatial distributions of 7,519 internal phylogenetic branches. Following established protocols [41], a phylogenetic branch was considered present in a grid cell if any of its descendant species occurred there.

We applied two complementary algorithms to address distinct conservation objectives. For regional conservation objective, we used the Core Area Zonation (CAZ2) algorithm, which emphasizes complementarity to build an efficient, representative network that retains core areas for all features, particularly rare taxa. For local conservation objective, we used the Additive Benefit Function (ABF), which prioritizes areas of high local biodiversity value.

Furthermore, we implemented two weighting regimes for these biodiversity features. Under uniform weighting, all species and phylogenetic branches received equal weight (1.0). Under conservation-priority weighting, individual species weights were incremented based on their threat status [42] (China’s Biodiversity Red List: Higher Plants), legal protection [43] (China’s National Key Protected Plants List), and QTP endemism [44]. For phylogenetic features, to properly integrate evolutionary history, the weight of each internal branch was set equal to its branch length. This ensured that long, distinctive evolutionary lineages contributed disproportionately to the overall conservation value [6,41].

### 2.7 Performance of growth forms as conservation surrogates

To evaluate the performance of growth forms as conservation surrogates, we delineated priority conservation areas (PCAs) under two global targets: the‘30×30’target [45] (protecting 30% of the land area, corresponding to grid cells with a Zonation rank ≥ 0.7) and the‘Half-Earth’target [46] (protecting 50% of the land area, rank ≥ 0.5).We then quantified the spatial similarity—using the Jaccard similarity index—between the PCAs derived for all plants and for each of the five growth forms, across the two conservation objectives, weighting regimes, and protection targets.

Next, to assess the reciprocal effectiveness of the all plants and individual growth forms as conservation surrogates, we employed a dual quantitative framework following established protocols [41]. For the regional conservation objective (CAZ2 algorithm), effectiveness was quantified by measuring the proportion of total taxonomic (TD) and phylogenetic (PD) diversity retained as potential conservation areas were sequentially expanded according to priority rankings (Supplementary Method 3, Equations S1 and S2). A species or phylogenetic branch was considered adequately represented when at least 30% of its distribution range was included within the PCAs, aligning with the representation targets of the Global Biodiversity Framework [45]. For the local conservation objective (ABF algorithm), prioritization efficacy was quantified using two metrics emphasizing restricted-range features: Weighted Endemism (WE) and Phylogenetic Endemism (PE), calculated based on the cumulative representation of endemic species and endemic phylogenetic branch lengths, respectively (Supplementary Method 3, Equations S3 and S4).

Finally, to systematically identify the most cost-effective surrogates, we formulated a novel efficiency metric: the Ability of Conservation Indicator (ACI). Anchored to the Global Strategy for Plant Conservation [47], we adopted its stringent target of capturing 75% of known diversity as our performance benchmark. The ACI evaluates the spatial economy of a surrogate by comparing the minimum land area required to meet this target using the target group directly against the area required when using a surrogate:

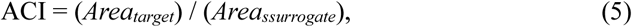

where *Area_target_* is the minimum proportion of the QTP area required to adequately represent 75% of the target group’s diversity (TD or PD) by prioritizing the target group directly, and *Area_surrogate_* is the corresponding proportion required to achieve the same 75% representation goal using a specific growth form as a surrogate. A higher ACI value denotes greater relative efficiency. Specifically, an ACI value ≥ 1 indicates a highly efficient surrogate that equals or outperforms the direct prioritization of the aggregate target group. This counter-intuitive result occurs when a specific growth form acts as a strong environmental filter for high-quality habitats, guiding conservation actions to key biodiversity areas more rapidly than the spatially diffuse all plants aggregate.

## 3. Results

### 3.1 Divergent spatial patterns across plant growth forms

Pronounced disparities in geographical range size among the five growth forms were paralleled by contrasting spatial diversity patterns. Herbs occupied the most extensive range (2.58 × 10⁶ km², covering 93.9% of the QTP), exhibiting a near-complete plateau-wide distribution that mirrored the pattern of the all plants. By contrast, trees, climbers, and epiphytes were virtually absent from the Plateau Platform, remaining restricted primarily to the Himalaya and Hengduan mountains (Table 1; Fig. 2; Fig. S1).

**Table 1.** Geographical range size (km²) and proportional area coverage (%) of the aggregate vascular flora (All plants) and each plant growth form across the Qinghai–Tibet Plateau (QTP). Range size is defined as the cumulative area of all grid cells where the species richness of the corresponding group exceeds zero (i.e., the total area occupied by at least one species).

| Growth forms | Range size (km<sup>2</sup>) | Proportion of QTP (%) |
| --- | --- | --- |
| All plants | 2.58×10<sup>6</sup> | 93.9 |
| Climbers | 1.11×10<sup>6</sup> | 40.5 |
| Epiphytes | 0.63×10<sup>6</sup> | 22.9 |
| Herbs | 2.58×10<sup>6</sup> | 93.8 |
| Shrubs | 1.76×10<sup>6</sup> | 64.1 |
| Trees | 1.26×10<sup>6</sup> | 45.7 |

**Fig. 2.**
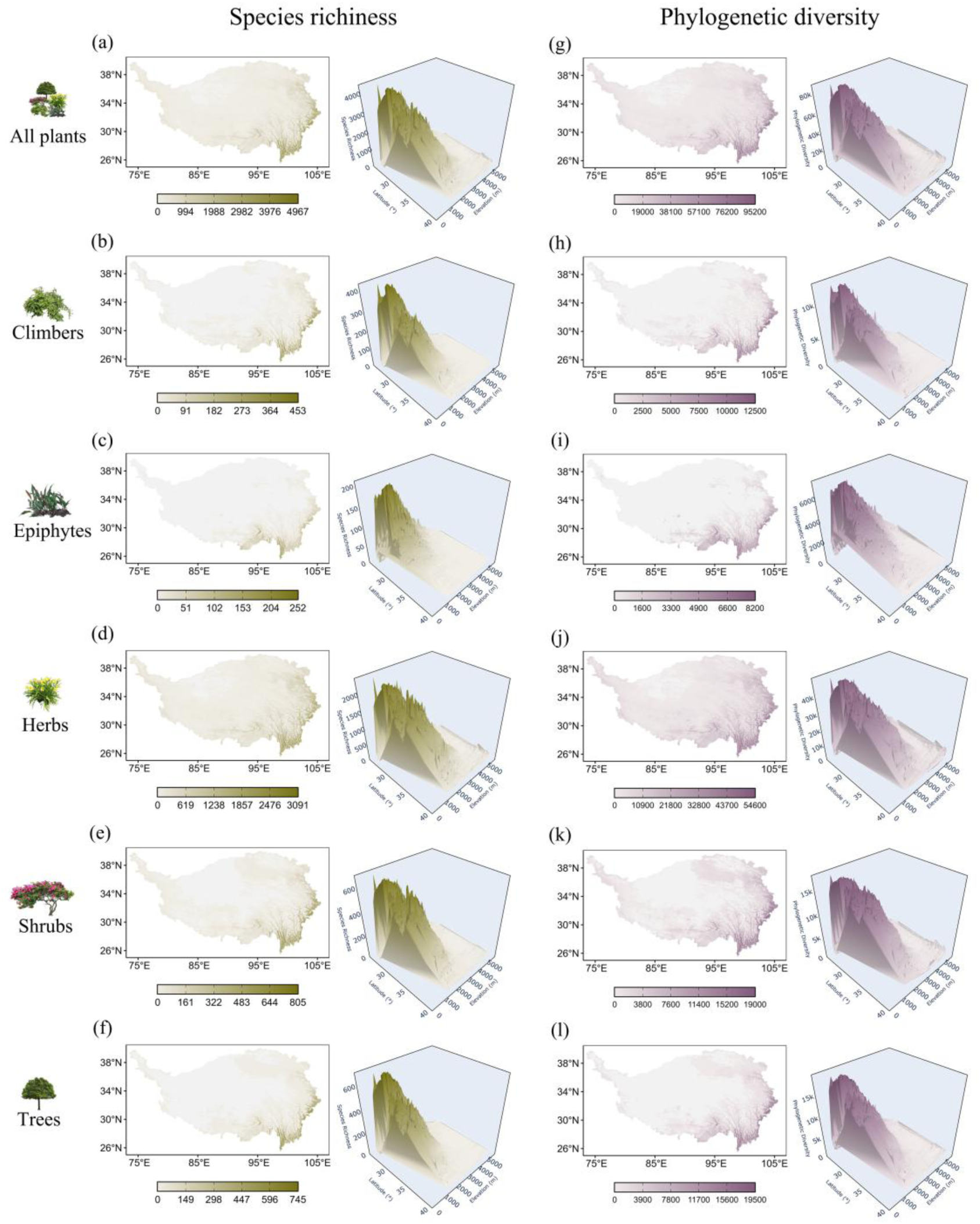
Divergent latitudinal and elevational gradients drive the decoupling of taxonomic and phylogenetic diversity across growth forms. (a–f) Geographic patterns of Species Richness (SR) for the all plants (a) and five major growth forms: climbers (b), epiphytes (c), herbs (d), shrubs (e), and trees (f). (g–l) Corresponding patterns of Faith’s Phylogenetic Diversity (PD; defined as the total branch length of the minimal subtree connecting all species in an assemblage). Panels display the all plants (g) followed by the corresponding growth forms: climbers (h), epiphytes (i), herbs (j), shrubs (k), and trees (l). For both metrics, inset trend surfaces illustrate the joint effects of latitude and elevation, revealing divergent environmental responses across growth forms; statistical details are provided in Table S4. Color bars apply to both the map and the trend surface within each panel.

Spatial correlation analyses confirmed these divergences. The spatial congruence between herbs and all plants was near-perfect (Spearman’s r > 0.99, P < 0.001; Fig. S2). Conversely, epiphytes exhibited the most distinct spatial constraints, showing the weakest association with other growth forms, particularly herbs (r = 0.66–0.70, P < 0.001).

Diversity metrics (SR, PD, WE, and PE) for all plants, herbs, and shrubs generally followed a southeast-to-northwest decline. However, the western QTP emerged as a notable exception: relatively high WE and PE values in this region revealed localized centers of endemism that diverged from overall richness patterns (Fig. 2). Meanwhile, the diversity of trees, climbers, and epiphytes was highly concentrated in the lower latitudes and elevations of the southeastern and eastern QTP. Along environmental gradients, these metrics displayed divergent responses. Both SR and PD exhibited unimodal (hump-shaped) relationships with elevation, yet the elevation of peak diversity shifted systematically across growth forms—from ∼3,300 m for herbs down to ∼2,200 m for epiphytes (Fig. 2; Fig. S3 and Table 4). Latitudinally, SR and PD increased toward the warmer tropics, peaking around 28–29°N. In contrast, neither WE nor PE exhibited clear linear or unimodal relationships with elevation or latitude (Figs. S3–S4).

### 3.2 Growth forms composition of plant assemblages

The QTP’s plant assemblages across the QTP were characterized by the overwhelming dominance of herbs, constituting 62.7% of total taxonomic diversity (TD) and 57.8% of phylogenetic diversity (PD) (Fig. S5a). While this dominance persisted across major physiographic units, it varied systematically with geography (Fig. S5b). Specifically, herb prevalence intensified toward the cold, arid northwest—exceeding 80% in alpine steppes and meadows where non-herb forms were virtually absent. In contrast, a structural shift occurred in forest-dominated sub-ecoregions: the monopoly of herbs diluted, yielding a more balanced distribution of growth forms, a pattern mirrored in the phylogenetic composition (Fig. S6). Grid-cell-level analyses further quantified this ubiquity, with herbs accounting for 89.0 ± 0.8% (mean ± SE) of TD and 81.0 ± 1.2% of PD locally. Conversely, substantial contributions from non-herb forms remained spatially constrained to the Eastern Himalayas, the southern Hengduan Mountains, and the eastern QTP margins (Fig. 3).

**Fig. 3.**
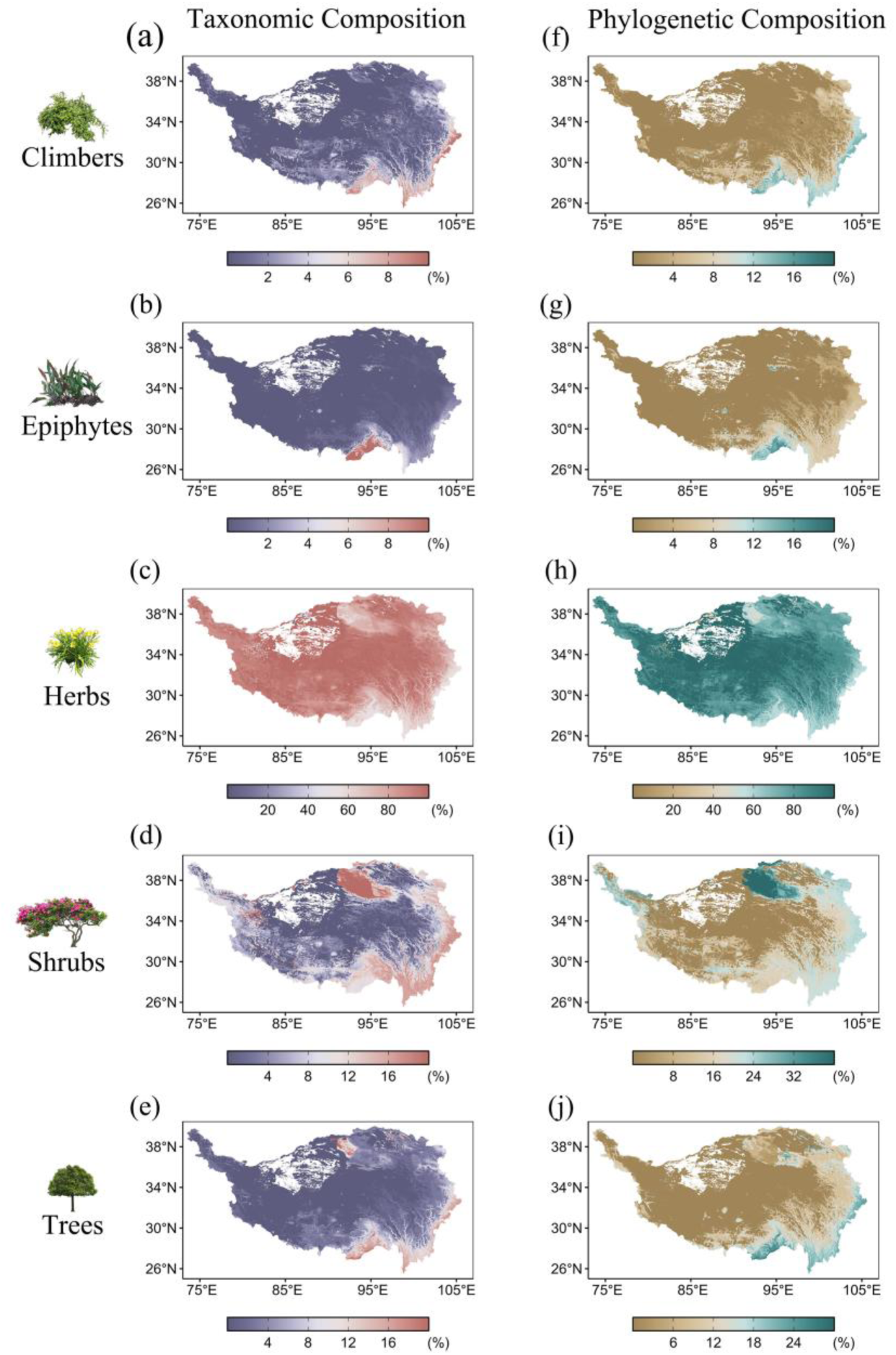
Taxonomic and phylogenetic composition of growth forms within Qinghai–Tibet Plateau plant assemblages. (a–e) Spatial variation in the taxonomic contribution of plant growth forms, quantified as the proportion of total species richness represented by each group within a 1-km² grid cell. Panels show the relative contributions of climbers (a), epiphytes (b), herbs (c), shrubs (d), and trees (e). (f–j) Corresponding phylogenetic contribution, expressed as the proportion of total assemblage-level phylogenetic diversity (PD) captured by species within each growth form. Panels follow the same growth-form order as in (a–e). Color bars indicate the proportional contribution (0–100%) for taxonomic (a–e) and phylogenetic (f–j) dimensions, respectively.

### 3.3 Effectiveness of growth forms as conservation surrogates

Priority conservation areas (PCAs) for both TD and PD were consistently concentrated in the southeastern and eastern QTP (Figs. S7–S10), regardless of the specified conservation targets (30% vs. 50%) or spatial scales (regional vs. local). Despite this general trend, spatial similarity among individual growth forms remained notably low. In particular, epiphytes displayed the lowest spatial congruence with all other groups (Fig. 4; Figs. S11–S13).

**Fig. 4.**
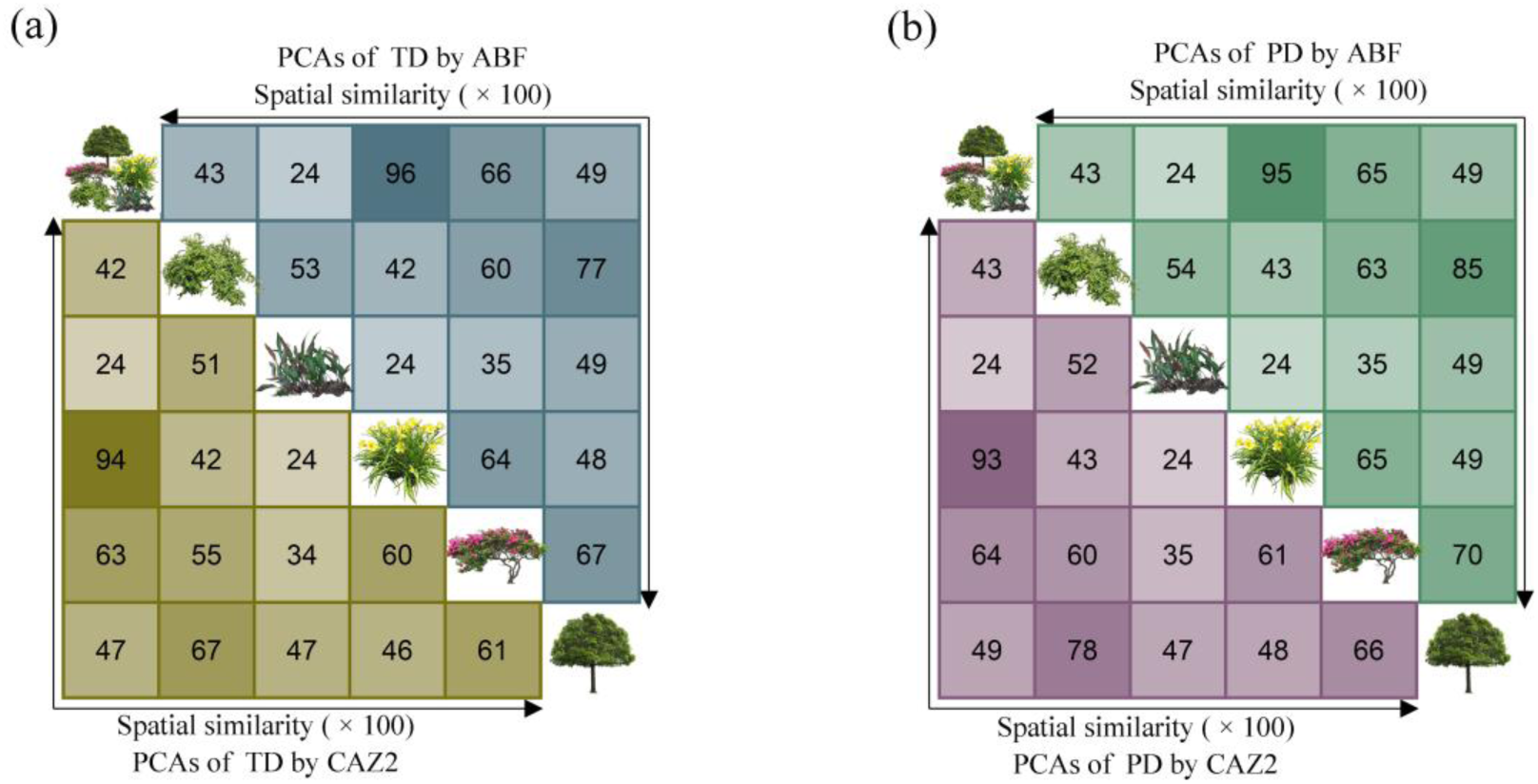
Spatial similarity among priority conservation areas across diversity facets and prioritization algorithms on the Qinghai–Tibet Plateau. Priority conservation areas (PCAs) were identified for all vascular plants and for each of the five plant growth forms under a 30% area protection target, using a weighted prioritization scheme implemented in Zonation 5. Two algorithms were compared: Core Area Zonation (CAZ2; prioritizing regional complementarity) and Additive Benefit Function (ABF; prioritizing local representativeness). (a) Spatial similarity based on Taxonomic Diversity (TD). The matrix displays pairwise Jaccard similarity indices between growth forms: the lower-left triangle shows incongruence using the CAZ2 algorithm, while the upper-right triangle shows incongruence using the ABF algorithm. (b) Spatial similarity based on Phylogenetic Diversity (PD), presented in the same matrix format as in (a). Darker colors indicate higher spatial similarity (higher Jaccard index), highlighting the degree of congruence between conservation priorities.

In potential conservation expansion simulations, the rate of diversity capture varied markedly across growth forms. Trees and climbers exhibited the steepest accumulation curves, indicating highly concentrated diversity, whereas herbs showed the shallowest curves, reflecting their diffuse distribution. Consequently, herbs consistently required the largest absolute land area to meet these conservation targets (Fig. 5a, b; Figs. S14–S16a, b).

**Fig. 5.**
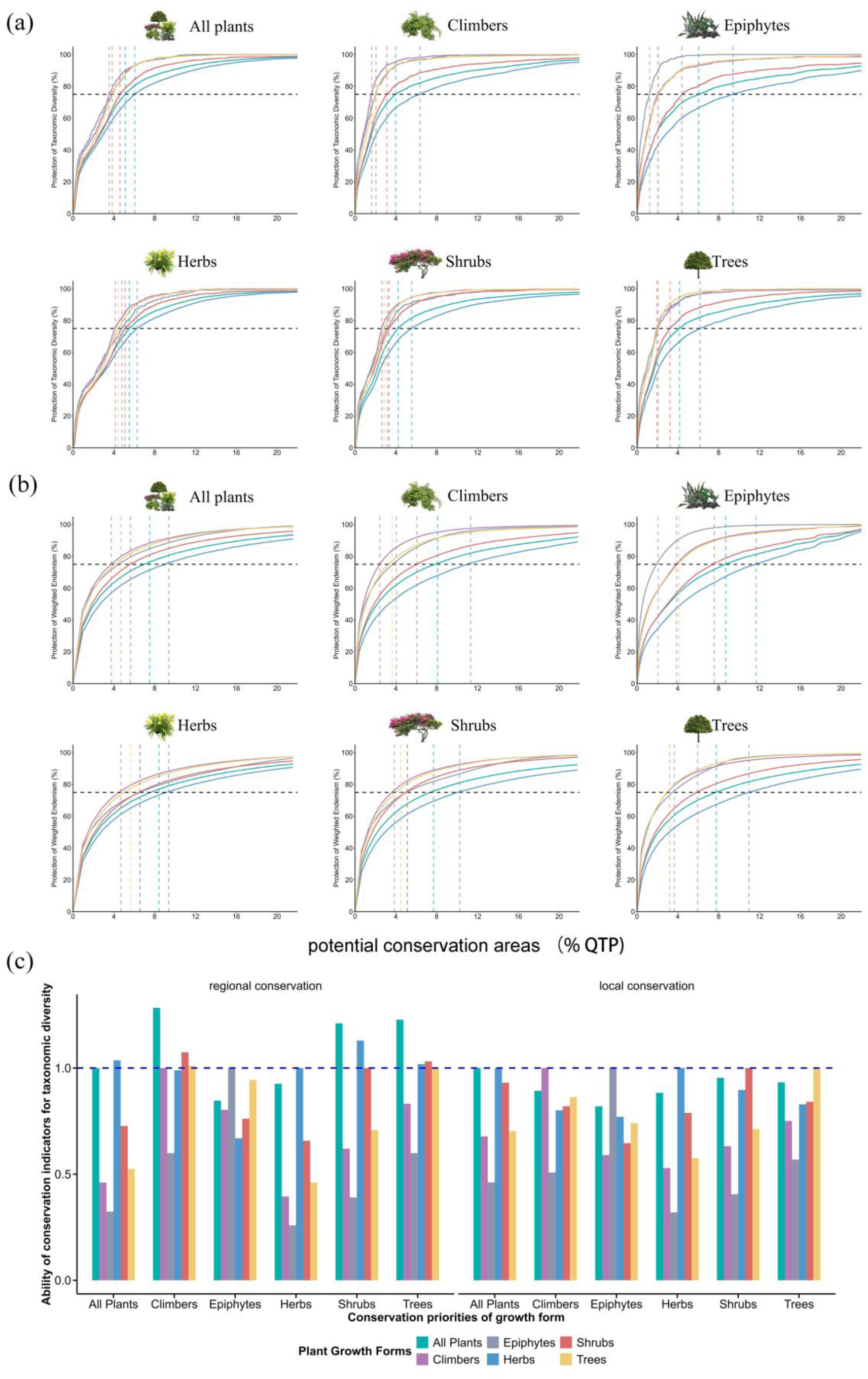
Assessment of conservation effectiveness and surrogacy performance across plant growth forms. (a–b) Cumulative diversity protection curves, evaluating how effectively conservation priorities identified for a specific growth form (surrogate) protect the broader diversity. Panels show the accumulation of (a) Regional Taxonomic Diversity (proportion of total species richness) and (b) Local Endemism (cumulative weighted endemism). Gray horizontal dashed lines indicate the 75% protection target; vertical dashed lines denote the minimum land area required to achieve this target. Note the steeper curves for trees/climbers compared to herbs. (c) Surrogacy performance, quantified by the Ability of Conservation Indicator (ACI). The blue dashed line (ACI = 1) represents performance equivalent to a specific prioritization for the target group itself. All analyses are color-coded: climbers (purple), epiphytes (grey), herbs (blue), shrubs (red), trees (yellow), and the aggregate flora (“all plants”; green). Conservation priorities were calculated using Zonation 5 with weighted prioritization schemes.

Crucially, we identified climbers and trees as the most effective umbrella growth forms. Evaluated by the Ability of Conservation Indicator (ACI) metric, climbers and trees outperformed all plants composite as robust surrogates for most other growth forms (TD ACI: 0.99–1.28 and 0.83–1.23, respectively). This preeminent role of climbers and trees was robustly confirmed from an evolutionary perspective, as they consistently yielded the highest ACI values for capturing the PD and phylogenetic endemism (PE) of other growth forms (Fig. 5c; Figs. S14–S16c).

## 4. Discussion

Understanding the spatial distribution of multifaceted biodiversity across plant growth forms is critical for effective conservation planning [10,48]. Our analysis across the QTP reveals pronounced spatial incongruencies in taxonomic and phylogenetic diversity patterns among growth forms. Consequently, conservation priorities predicated on aggregate metrics of total plant diversity—which are inherently skewed toward dominant groups—risk overlooking the evolutionary heritage of less abundant but functionally critical lineages. We identify trees and climbers as keystone functional groups that act as highly effective umbrella growth forms. Their distributions efficiently capture the broader taxonomic and phylogenetic diversity of other forms. By demonstrating that specific structural architects can robustly anchor conservation planning, we provide a tractable, trait-informed framework to shift the focus from individual species to critical functional dimensions.

### 4,1 Disparate diversity patterns across plant growth forms

The pronounced disparities in geographic range size across plant growth forms (Table 1) reflect a fundamental life-history strategy continuum [8,15,48]. Herbs epitomize a rapid-colonization strategy: their short generation times and versatile dispersal capacities enabled them to rapidly exploit the plateau’s heterogeneous environments, capitalizing on post-glacial expansion opportunities [11,18,49]. Conversely, the confinement of trees, climbers, and epiphytes to the southeastern margins is dictated by a strict ’double filter’: a primary ecophysiological barrier imposed by extreme cold and altitude beyond the treeline (∼4,900 m) [9,14,50], compounded by their intrinsically limited dispersal capabilities and reliance on specific microhabitats [8,48].

These intrinsic differences in niche breadth and dispersal capacity fundamentally decouple spatial biodiversity patterns among growth forms [17,48]. Crucially, we demonstrate that environmental filtering operates in a highly growth-form-specific manner, as evidenced by the systematic divergence in their elevational optima (Fig. S3). This marked incongruence challenges the primacy of aggregate diversity metrics—which are heavily skewed by dominant herbaceous taxa—as universal proxies for all functional groups [10–11]. Extrapolating biodiversity patterns from dominant clades to distinct functional components is therefore systematically flawed. Ultimately, our findings underscore the absolute necessity of a growth-form-explicit framework to accurately resolve community assembly mechanisms and effectively safeguard the full spectrum of mountain plant diversity [9].

### 4.2 The dominant role of herbs within plant assemblages

Moving beyond simple taxonomic inventories, our study highlights the functional composition of plant assemblages—a fundamental driver of ecosystem functioning [9]. Our macroecological synthesis demonstrates the overwhelming dominance of herbs across the QTP flora, both in terms of species richness and phylogenetic lineage accumulation (Fig. 3; Figs. S5-S6). This primacy is deeply rooted in evolutionary history. Since the late Miocene, the Hengduan Mountains have functioned as a formidable ’species pump,’ driving explosive radiations, particularly within herbaceous clades. The subsequent extinction of paleo-woody flora, coupled with an influx of cold-adapted herbs, firmly cemented this herbaceous dominance [3,19,51]. This shared biogeographic legacy [19,49,51] underpins the remarkable structural consistency observed across the QTP’s major physiographic units (Fig. S5b). Notably, the relatively recent, rapid diversification of these herbs (∼7–8 Ma) [19, 51] may explain their lower contribution to phylogenetic diversity compared to their taxonomic richness (Fig. 3; Figs. S5-S6).

Beneath this broad-scale uniformity lies pronounced scale-dependent heterogeneity driven by a distinct spatial gradient. As one moves from the species-rich southeastern margins toward the cold, arid northwest, herb dominance intensifies dramatically. Consequently, the proportional abundance of herbs is negatively correlated with overall plant diversity (Table S5). Within the southeastern tropical and subtropical forests—the QTP’s primary biodiversity hotspots—the herbaceous fraction drops to 50 – 60%, mirroring the structural evenness of tropical ecosystems [9,14]. Peak biodiversity in these assemblages is therefore maintained not by the monopolization of a single growth form, but through enhanced niche complementarity across diverse functional strategies [9,52]. Furthermore, because trees do not constitute the majority of species richness even in these structurally complex forests [53], our findings directly challenge prevailing tree-centric biases in biodiversity monitoring. This exposes an urgent need to shift conservation and restoration metrics away from simple afforestation targets toward a holistic, whole-community perspective [9,11,13].

### 4.3 Effectiveness of plant growth forms as conservation indicators

Our findings expose a fundamental conservation paradox across the QTP: herbs, despite overwhelmingly dominating regional assemblages, are inadequate surrogates for safeguarding overall functional diversity. Because priority conservation areas for herbs exhibit low spatial similarity with those of other growth forms (Fig. 4; Figs. S11–S13), strategies targeting this dominant group in isolation capture multidimensional diversity highly inefficiently (Fig. 5a, b; Figs. S14–S16a, b). This reinforces the emerging consensus that conservation planning predicated on taxonomically pooled data effectively masks functional identity and compromises multifaceted biodiversity targets [6,10–11].

To overcome the limitations of these simplistic proxies, we identify trees and climbers as highly effective umbrella growth forms. As foundational architects of ecosystem organization, these structural keystones deliver disproportionate conservation leverage; their prioritization yields superior cross-taxon efficacy that systematically safeguards the vast majority of other growth forms (Fig. 5c; Figs. S14–S16c). This surrogate power stems from their synergistic role in generating vertically complex forest architecture, which creates heterogeneous microhabitats, buffers against environmental extremes, and facilitates the coexistence of evolutionarily distinct lineages [9,52]. Within this structural framework, climbers—historically marginalized in conservation—function not as mere appendages, but as vital interstitial connectors that amplify resource availability [5,14,24]. Consequently, protecting forests rich in these architectural cores inherently secures the diffuse spatial patterns of the herbaceous understory, resolving a spatial complexity that would otherwise confound regional conservation planning.

### 4.4 Conservation Implications

Operationalizing our findings requires a paradigm shift in spatial conservation planning across the QTP. Rather than relying on aggregate taxonomic surrogates, which disproportionately reflect herbaceous distributions, conservation frameworks must explicitly pivot toward structural keystones: trees and climbers. Embedding this combined structural indicator into spatial planning allows policymakers to maximize the conservation return on investment across all growth forms [54–55]. This approach provides a tangible, verifiable criterion for designating Key Biodiversity Areas (KBAs) and Other Effective Area-based Conservation Measures (OECMs) under the Kunming-Montreal Global Biodiversity Framework’s “30×30” target [56]. Protecting these structurally complex forests thus delivers dual benefits: safeguarding a disproportionate share of evolutionary heritage while simultaneously enhancing vital ecosystem services such as carbon sequestration [52,56].

Translating this structural paradigm from a conceptual ideal into a measurable policy metric requires overcoming historical management biases. Traditional forestry and conservation practices have remained predominantly tree-centric, often neglecting the critical ecological functions of climbers and understory flora [11,14,24]. Future monitoring frameworks must therefore explicitly quantify vertical structural complexity across diverse forest types and environmental gradients [9,14]. The integration of high-resolution remote sensing, airborne LiDAR, and advances in functional biogeography offers a robust technological pathway to achieve this [52,56,57]. Together, these tools can operationalize structural complexity as a scalable, dynamic metric for large-scale conservation monitoring.

Beyond structurally complex forests, securing the QTP’s full evolutionary heritage mandates tailored, growth-form-specific management across non-forested biomes. For widespread herbs and shrubs, conservation efforts must prioritize structurally simplified ecosystems—such as the Qaidam Basin, the Three Rivers Source Region, and the Pamir-Western Kunlun Mountains—where these communities exhibit acute vulnerability to climate change and environmental degradation [58]. Within existing national parks, ecological resilience should be bolstered through targeted interventions, including seed bank augmentation for endemic herbs and browsing control for vulnerable shrubs, guided by integrated field and remote-sensing networks. Furthermore, epiphytes—the most range-restricted and biogeographically distinct growth form—face imminent threats from primary forest degradation and large-scale infrastructure development [17]. Their survival hinges on the uncompromising protection of remaining old-growth host-tree habitats and dedicated monitoring of their population dynamics in response to global change drivers.

### 4.5 Limitation and future directions

While growth-form identity provides a robust framework for conservation prioritization, operationalizing this approach requires addressing key methodological constraints. First, spatial sampling bias remains a persistent challenge in the macroecology of remote, hyper-diverse regions. In our dataset, 44.3% of species possess fewer than five occurrence records. This data deficit, typical of highly endemic montane floras, highlights an urgent need for targeted field surveys in chronically under-sampled frontiers, such as the Qiangtang Plateau and the Nyainqêntanglha–Gangdise transition zone. Such efforts are essential to ground-truth and iteratively refine the distribution models underpinning our priority maps (Figs. S7–S10). Second, future research must empirically extend this framework by integrating continuous functional traits (e.g., specific leaf area, wood density) [10]. This will determine whether discrete growth-form categorizations sufficiently capture multivariate functional strategies, or if continuous trait-based models reveal deeper mechanistic nuances. Furthermore, incorporating species abundance or biomass data [59] would shift the analytical focus from binary presence/absence to ecological dominance and functional impact—distinguishing, for instance, the disproportionate carbon and structural contributions of the giant *Cupressus torulosa* from the diminutive herb *Spiranthes sinensis*—thereby enabling conservation strategies that directly target ecosystem processes.

Ultimately, our findings demonstrate how growth-form-explicit patterns can refine spatial conservation priorities. We therefore propose a testable hypothesis for the global conservation community: future protected-area expansion on the QTP—if explicitly guided by priorities that maximize structural complexity (i.e., focusing on forests rich in trees and climbers)—will secure significantly greater overall biodiversity and ecosystem functionality than expansion driven solely by traditional species-richness hotspots. Testing this proposition addresses a core debate in conservation science: whether structural complexity should be elevated to a primary planning criterion alongside taxonomic diversity. A positive empirical outcome would not only validate our framework for the QTP but also establish a highly transferable model for safeguarding topographically complex, data-limited ecosystems worldwide.

## 5. Conclusion

Plant growth form acts as a fundamental driver of spatial biodiversity patterns across the QTP. Our analysis reveals that traditional, species-centric conservation planning—which heavily reflects dominant herbaceous taxa—is structurally blind and highly inefficient for protecting multidimensional biodiversity [54–55].

By demonstrating that trees and climbers function as superior umbrella growth forms, we provide an evidence-based blueprint for maximizing protection efficacy. Moving forward, explicitly integrating growth-form identity and forest structural complexity into spatial planning offers not only a quantitative foundation for preserving the QTP’s unique flora but also a highly transferable, scalable strategy to enhance conservation outcomes in global biodiversity hotspots.

## Data availability statement

All data needed to evaluate the conclusions in the paper are present in the paper and/or the Supplemental Information. Species occurrence data are available from the Global Biodiversity Information Facility (GBIF) at (https://www.gbif.org/occurrence/download/0049768-241126133413365), Additional herbarium and field data are available from the lead contact upon request. Climate data are publicly available at https://chelsa-climate.org/. Topographic data are available at https://www.earthenv.org/DEM. Soil data are available from the Harmonized World Soil Database v2.0 at https://www.fao.org/soils-portal/data-hub/soil-maps-and-databases/harmonized-world-soil-database-v20/en/.

## Acknowledgements

The authors thank Liang Hu for providing the checklist of climbing plants in China. Funding for this research came from the Second Tibetan Plateau Scientific Expedition and Research Program (Grant No. 2019QZKK0502) and the National Natural Science Foundation of China-Yunnan Joint Fund (Grant No. U25A20637).

## Author contributions

Feng Liu, Zhiyuan Liang, Bo Li, and Jinming Hu conceptualized the study. Feng Liu, Zhiyuan Liang, Jin Ye, Jie Li and Feiling Yang collected all data. Feng Liu, Zhiyuan Liang, Xiao Xu and Jingge Kuang conducted the investigation. Feng Liu, Zhiyuan Liang, Bo Li, Yupeng Geng and Jinming Hu wrote the manuscript. All authors have read and approved the final manuscript.

