## Supplementary method,Fig S1~S16, Table S1~S5 for "Plant growth forms shaping biodiversity patterns can guide conservation planning on the Qinghai-Tibet Plateau"

**Supplementary materials for**

### Plant growth forms govern biodiversity patterns and inform large-scale conservation planning on the Qinghai-Tibet Plateau

Feng Liu[1], Zhiyuan Liang[2], Xiao Xu[1], Jingge Kuang[1], Jin Ye[3], Filing Yang[3], Jie Li[3], Yupeng Geng[1], Bo Li[1 *], Jinming Hu[4]

### Appendix

**Supplementary method 1** a vascular plant checklist for the Qinghai-Tibet Plateau

Establishing a comprehensive and taxonomically consistent species checklist is a critical prerequisite for all subsequent analyses. To resolve discrepancies in estimates of the total vascular flora of the Qinghai-Tibet Plateau (QTP) [1-2], we constructed an integrated checklist by synthesizing data from multiple authoritative sources. These included: (1) core national and regional floras, such as Flora Reipublicae Popularis Sinicae, Flora of China, and The Vascular Plants and Their Eco-geographical Distribution of the Qinghai-Tibet Plateau [3]; (2) updated regional inventories, including the Catalogue of Plant Species in China [4] and recent regional checklists [5-7]; and (3) global taxonomic backbones, specifically the World Checklist of Vascular Plants [8], supplemented by recent peer-reviewed literature [1,9-10].

To ensure nomenclatural consistency, we standardized all species names primarily against the Catalogue of Plant Species in China [4], with further alignment against WCVP and the Catalogue of Life. This process involved automated name-matching followed by extensive manual verification. We retained only species-level taxa, excluding hybrids and aggregating infraspecific taxa to the species level. Our final, curated checklist comprises 17,844 vascular plant species (representing 2,462 genera and 271 families), forming the foundational dataset for this study.

**Supplementary method 2** Occurrences records cleaning

To mitigate pervasive issues of sampling bias, geolocation errors, and taxonomic inconsistencies in raw biodiversity databases, we implemented a rigorous, multi-stage cleaning protocol. First, we performed automated quality control using the CoordinateCleaner R package [11] to flag and remove: (i) fossil records; (ii) geographic outliers (e.g., records assigned to ocean coordinates or country centroids); (iii) duplicate entries; and (iv) records from biodiversity institutions (e.g., botanical gardens), which may not represent natural populations.

Subsequently, to ensure records reflected contemporary distributions with sufficient locational accuracy, we applied temporal and spatial filters. We excluded records collected before 1950 and those with a coordinate precision of fewer than three decimal places (implying an uncertainty of >100 m). Taxonomic standardization was conducted using the U.Taxonstand package [12 ] and cross-referenced against our curated checklist; all infraspecific taxa were aggregated to the species level.

Finally, to reduce spatial autocorrelation and sampling bias for robust SDM calibration, we applied spatial thinning. Using the spThin package [13], we retained only a single occurrence record within a minimum distance of 1 km for each species. This threshold matches the resolution of our environmental variables and effectively minimizes overrepresentation from densely sampled localities.

**Supplementary method 3** Detailed equations for Performance of growth forms as conservation surrogates

The reciprocal performance among all vascular plants and the five specific plant growth forms as conservation surrogates was evaluated based on two complementary conservation objectives. (1) For the regional conservation objective, we quantified surrogate performance by measuring the proportion of total taxonomic diversity (TD) and phylogenetic diversity (PD) captured as potential conservation areas were sequentially expanded according to prioritized rankings. For TD, surrogate effectiveness was defined as the cumulative proportion of species diversity protected relative to the total species pool (Equation S1).

$$TD(\%) = \frac{g}{G} \times 100, \qquad \text{(Equation S1)}$$

Where g denotes the number of protected species (at least 30% species range, consist with the 30 × 30 Global Biodiversity Framework target [14]), and G represents the total vascular plant species count on the QTP.

Phylogenetic performance was quantified as the percentage of total regional phylogenetic diversity (PD) captured within the simulated potential conservation areas, calculated from the branch lengths retained (Equation S2).

$$PD(\%) = \frac{\sum_{i=1}^{b} L_i}{\sum_{i=1}^{B} L_i} \times 100, \qquad \text{(Equation S2)}$$

Where *b* represents the number of protected phylogenetic branches (at least 30% of distribution range of internal phylogenetic branches), *B* denotes the total number of phylogenetic branches, and *L* indicates the length of branch *i*.

For the local conservation objective, the effectiveness of spatial prioritization was quantified using two metrics specifically targeting restricted-range biodiversity: Weighted Endemism (WE) and Phylogenetic Endemism (PE). WE was calculated as the sum of species richness weighted by the inverse of their range sizes, representing the concentration of rare taxa. Similarly, PE was employed to measure the retention of evolutionary history restricted to specific areas, calculated by weighting phylogenetic branch lengths by their geographic rarity.

$$WE(\%) = \frac{1}{N} \bullet \sum_{i=1}^{N} \left[ \frac{d_i}{D_i} \right] \times 100, \qquad \text{(Equation S3)}$$

Where N represents the total number of vascular plant species within a given growth form on the QTP, *d* denotes the portion of specie range overlapped by potential conservation areas for the species, and D indicates the total geographical range of the species.

$$PE(\%) = \frac{1}{\sum_{i=1}^{k} L_i} \bullet \sum_{i=1}^{k} \left[ L_i \bullet \frac{e_i}{E_i} \right] \times 100, \qquad \text{(Equation S4)}$$

Where k denotes the total number of branches of the tree, $e_i$ represents the spatial overlap between the distribution of the branch *i* and potential conservation areas, $E_i$ indicates the total distribution area of the branch *i*, and L is the length of branch *i*.

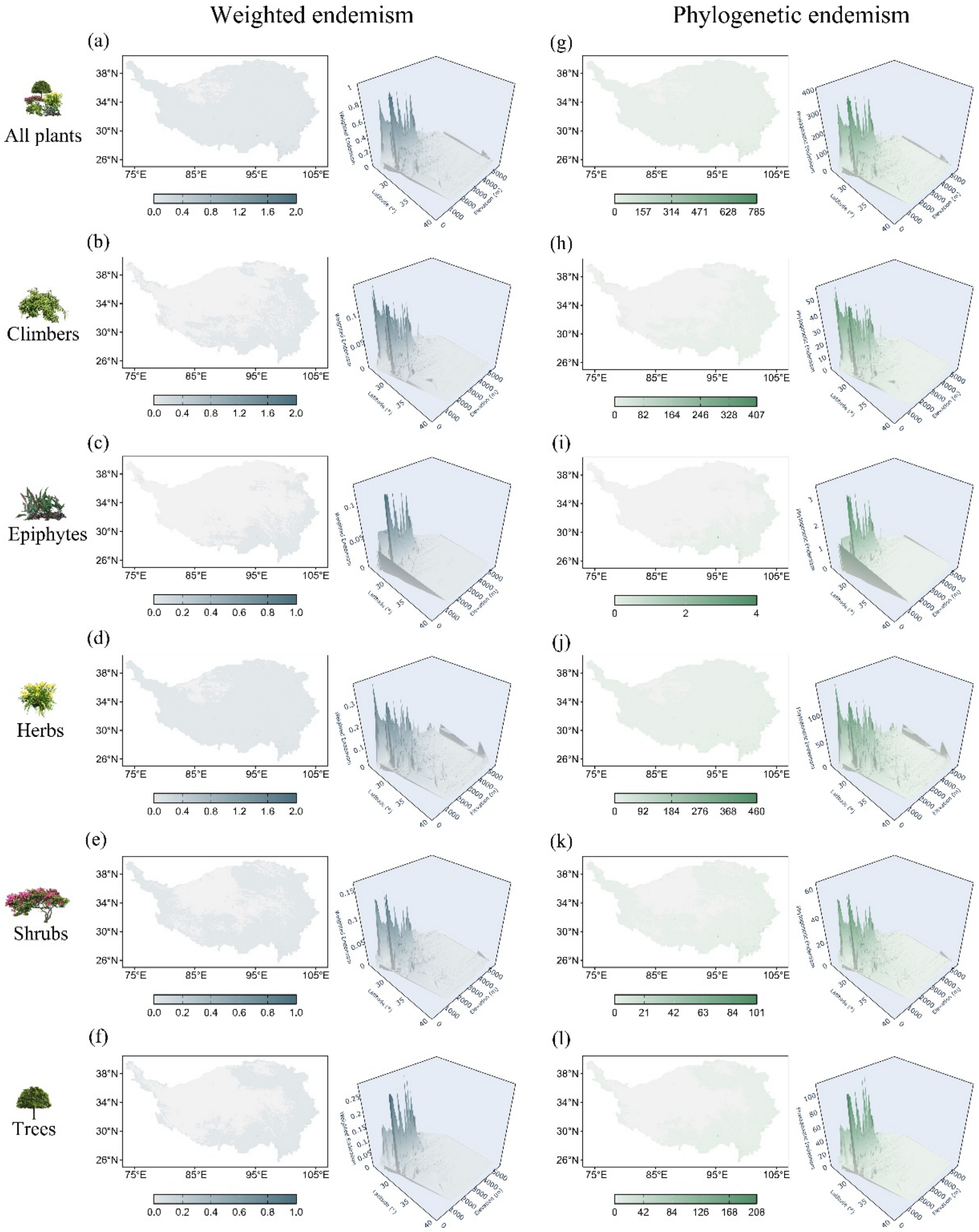


**Fig. S1. Latitudinal and elevational gradients of weighted and phylogenetic endemism for vascular plants across the Qinghai-Tibet Plateau.** (a–f) Geographic patterns of Weighted Endemism (WE), quantified as the sum of species richness inversely weighted by their range sizes, shown for all vascular plants (a) and five major growth forms: climbers (b), epiphytes (c), herbs (d), shrubs (e), and trees (f). (g–l) Corresponding patterns of Phylogenetic Endemism (PE), quantified as the sum of phylogenetic branch lengths inversely weighted by their spatial occupancy, shown for all plants (g) and individual growth forms (h–l). For both dimensions, the overlaid trend surfaces illustrate the joint effects of latitude and elevation; detailed statistical outputs are provided in Table S4. Color bars represent the value ranges for WE (a–f) and PE (g–l) respectively, and apply to both the geographic maps and their corresponding trend surfaces.

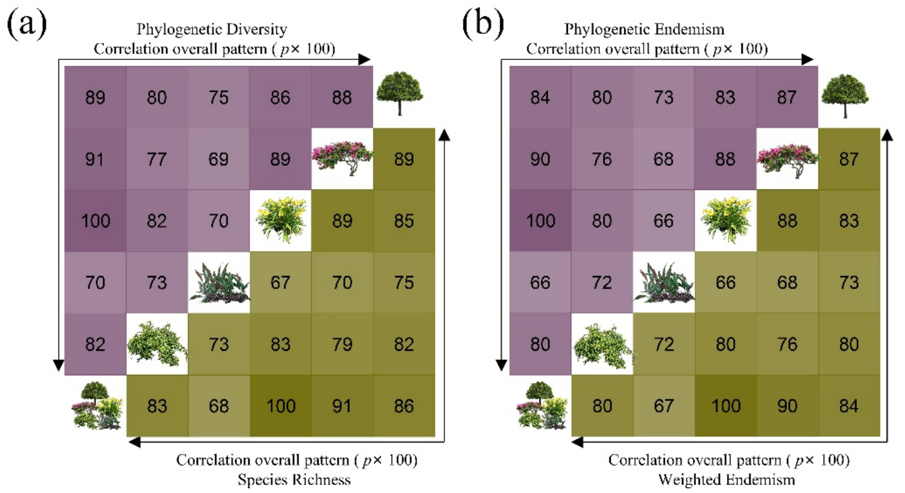


**Fig. S2. Spatial congruence in multi-faceted diversity patterns across plant growth forms.** Pairwise spatial congruence between diversity patterns of different growth forms and all vascular plants was quantified using Spearman's rank correlation coefficients. (a) Correlation matrix for Species Richness (lower triangle) and Phylogenetic Diversity (upper triangle). Each cell represents the strength of the spatial correlation between the corresponding groups for the respective metric. (b) Correlation matrix for Weighted Endemism (lower triangle) and Phylogenetic Endemism (upper triangle). In both matrices, the hierarchical order of rows and columns is: all vascular plants, climbers, epiphytes, herbs, shrubs, and trees. Color intensity indicates the magnitude of positive spatial correlation, with darker shades denoting higher congruence. All correlations shown are statistically significant ($p < 0.001) unless otherwise specified.

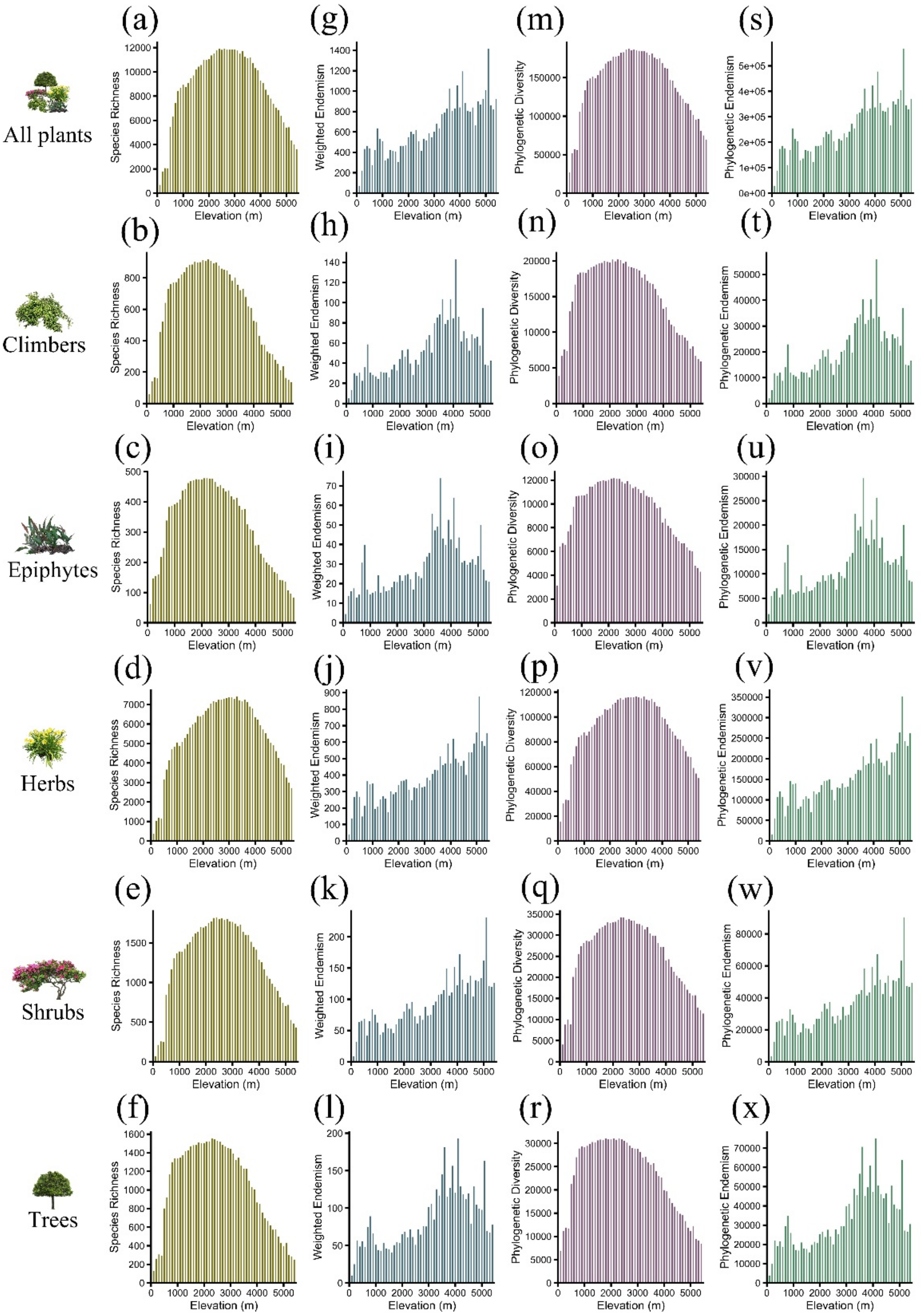


**Fig. S3. Elevational gradients of multi-faceted plant diversity across 100-m bands on the Qinghai-Tibet Plateau.** The figure displays the elevational distribution of four complementary diversity metrics for all vascular plants (all plants) and each of the five growth forms. Panels are organized by metric: (a–f) Species Richness, (g–l) Weighted Endemism, (m–r) Phylogenetic Diversity, and (s–x) Phylogenetic Endemism. Within each metric group, individual panels correspond to all plants, climbers, epiphytes, herbs, shrubs, and trees, respectively. All metrics were aggregated within 100-m vertical intervals to identify consistent or divergent patterns across life forms along the full elevational range of the Qinghai-Tibet Plateau.

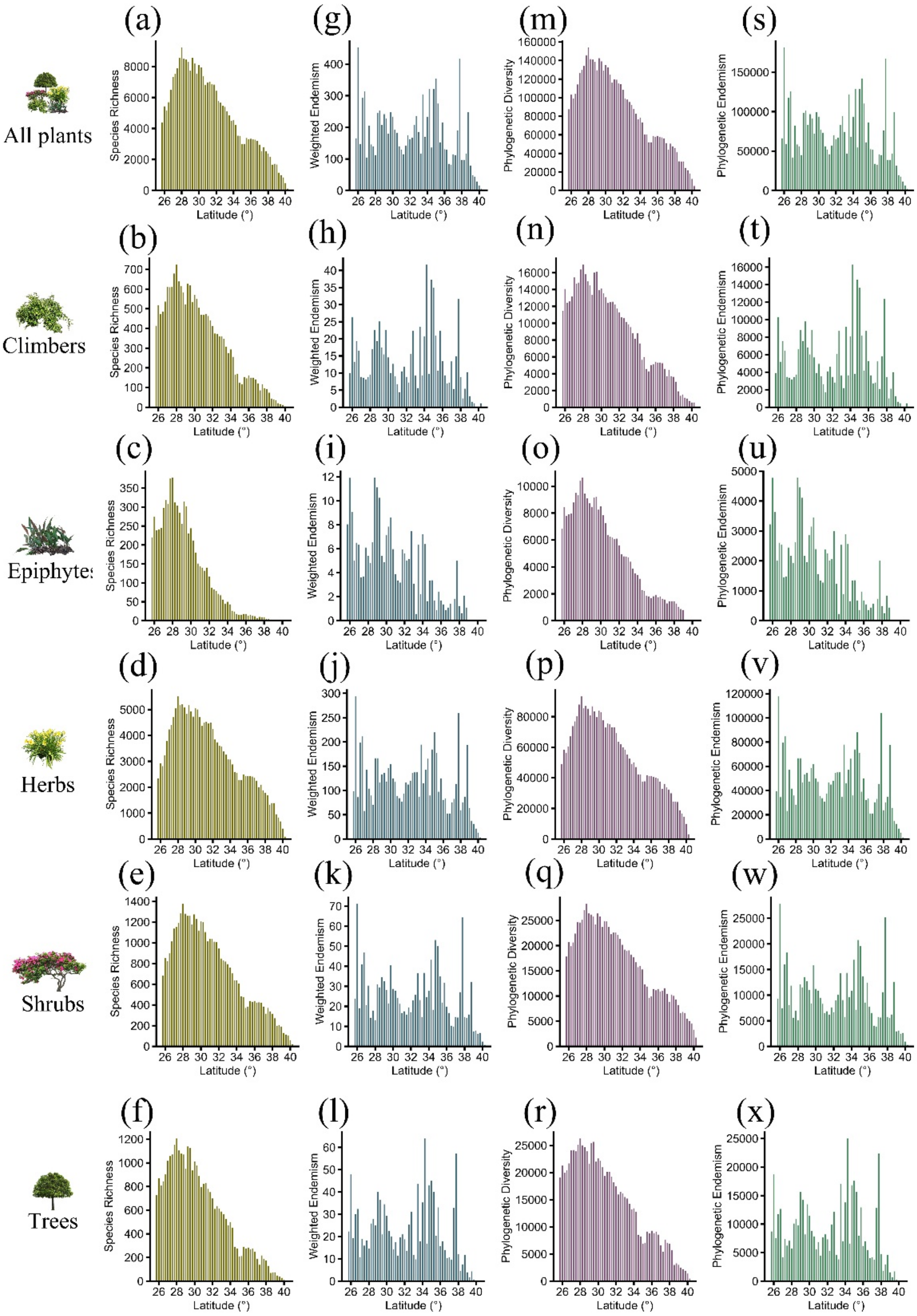


**Fig. S4. Latitudinal patterns of multi-faceted plant diversity across 0.25° bands on the Qinghai-Tibet Plateau.** The figure displays the latitudinal variation of four complementary diversity metrics for all vascular plants and individual growth forms across the Qinghai-Tibet Plateau. Panels are grouped by diversity dimension: (a–f) Species Richness, (g–l) Weighted Endemism, (m–r) Phylogenetic Diversity, and (s–x) Phylogenetic Endemism. Within each dimension, panels represent all vascular plants, climbers, epiphytes, herbs, shrubs, and trees, respectively.

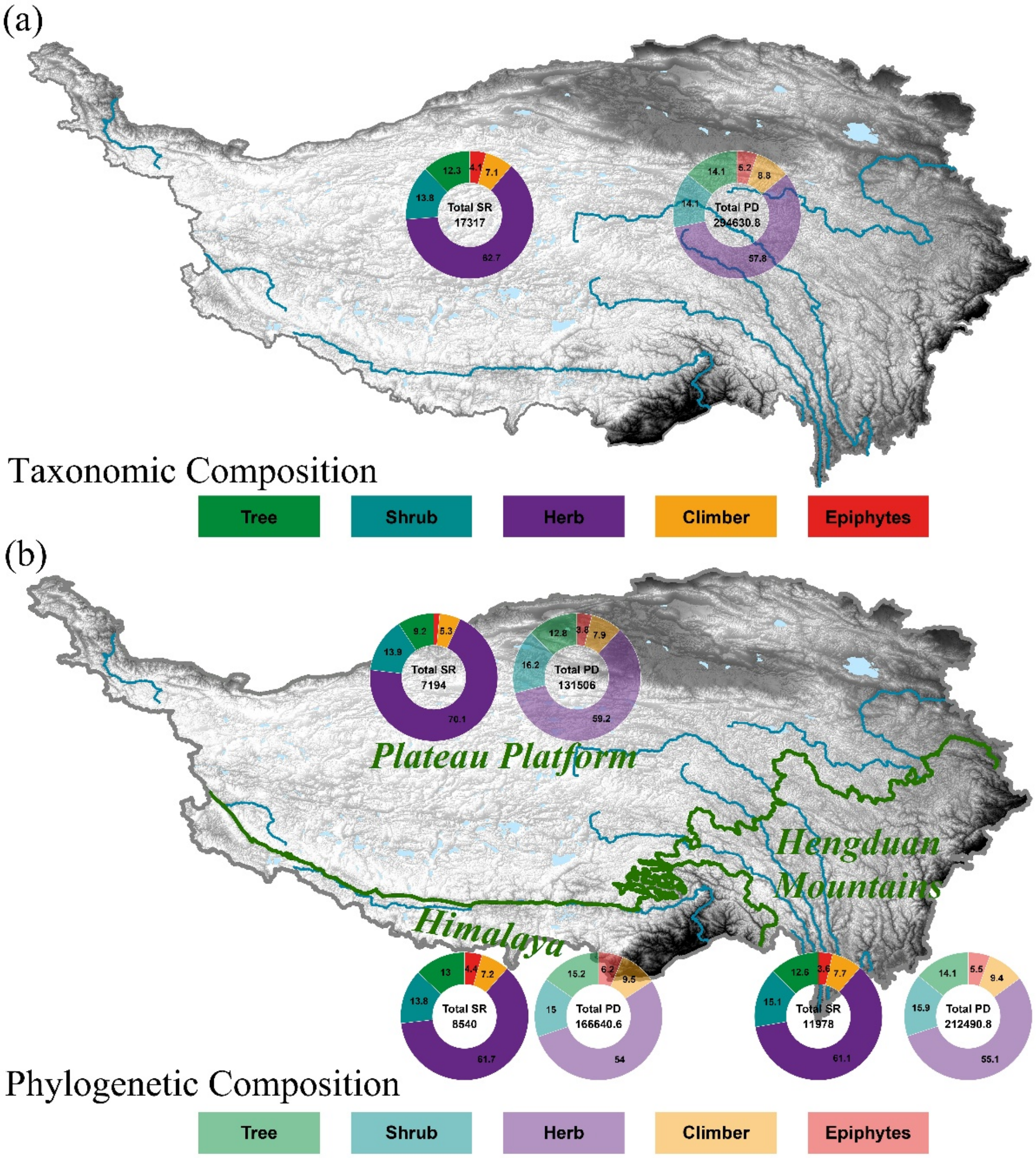


**Fig. S5. Composition of plant growth forms to taxonomic and phylogenetic diversity across the Qinghai-Tibet Plateau (QTP) and its major physiographic units.** (a) Overall composition of the vascular plant flora across the entire QTP. The ring segments with darker shades represent the proportion of total Species Richness (SR) contributed by each growth form, whereas segments with lighter shades indicate the proportion of total Phylogenetic Diversity (PD). (b) Comparative composition within three key biogeographic subregions: the Plateau Platform, the Himalayas, and the Hengduan Mountains. Growth forms are consistently color-coded across all panels: trees (yellow), shrubs (red), herbs (blue), climbers (purple), and epiphytes (grey). Note differences in the relative contribution of growth forms to SR versus PD, reflecting the evolutionary distinctiveness of specific lineages.

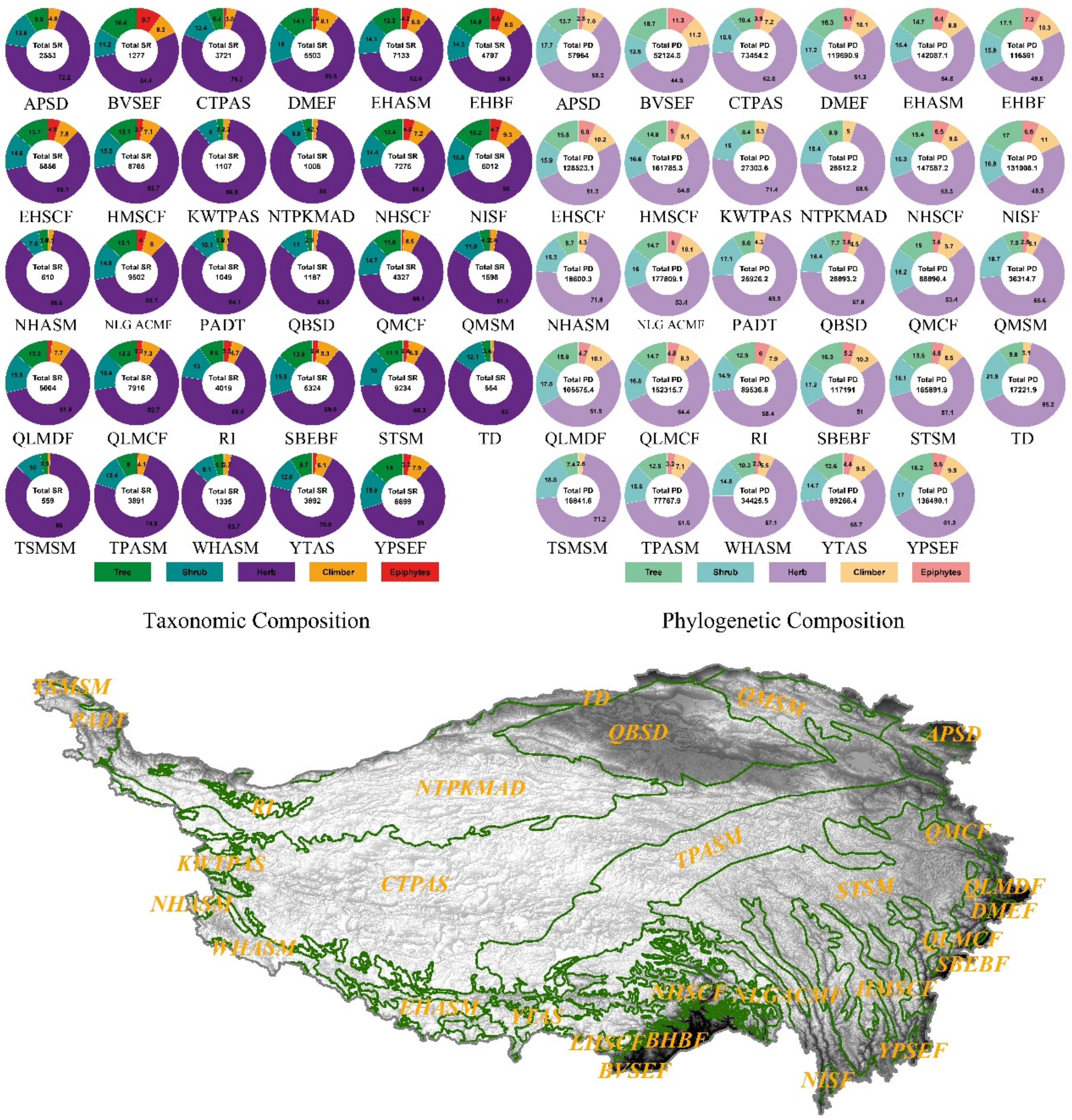


**Fig. S6. Spatial heterogeneity in plant growth form composition across the 29 sub-ecoregions of the Qinghai-Tibet Plateau.** The charts display the proportional contribution of each growth form to taxonomic and phylogenetic diversity within each distinct ecoregion. Similar to Supplementary Fig. 5, ring segments with darker shades represent the proportion of total Species Richness (SR), while segments with lighter shades indicate the proportion of total Phylogenetic Diversity (PD). Growth forms are consistently color-coded: trees (yellow), shrubs (red), herbs (blue), climbers (purple), and epiphytes (grey).

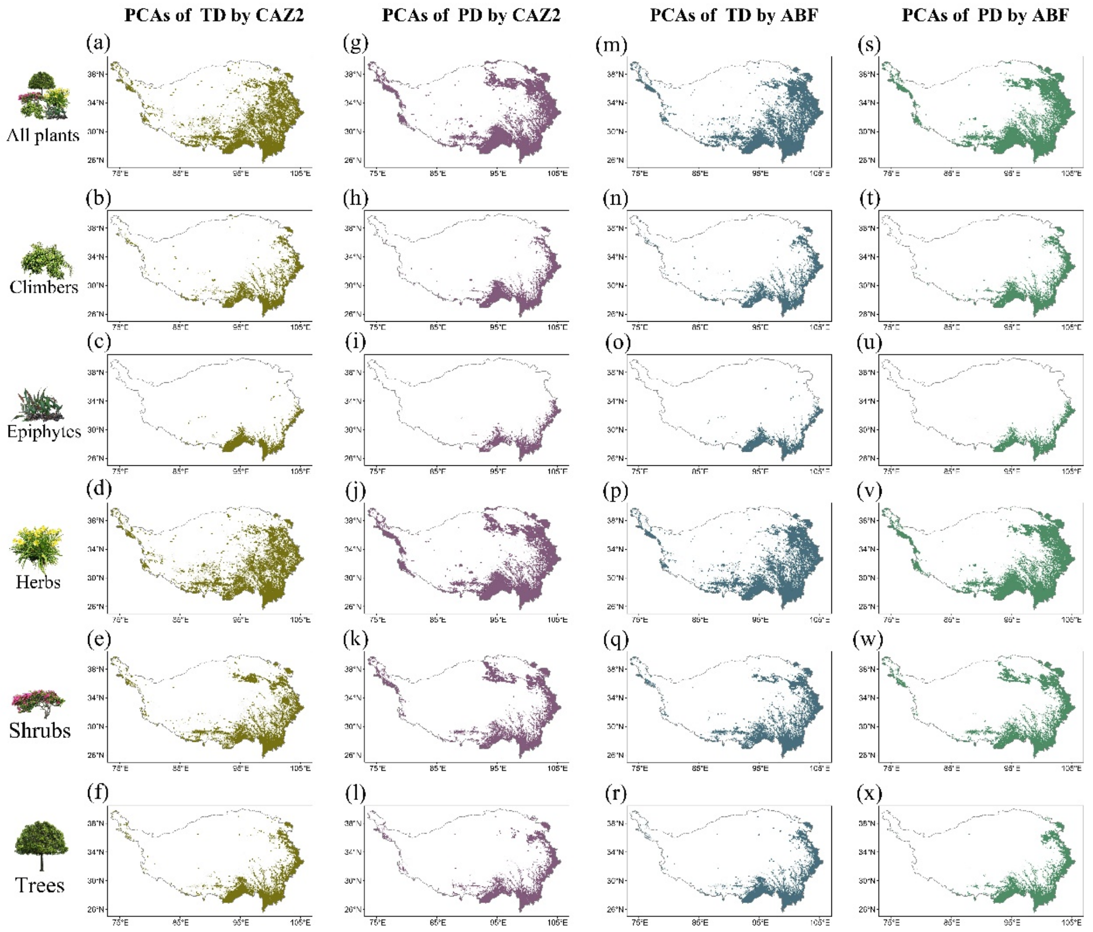


**Fig. S7. Spatial patterns of priority conservation areas (PCAs) across plant growth forms and diversity facets on the Qinghai-Tibet Plateau.** PCAs were identified for all vascular plants and each of the five growth forms under a 30% area-protection target, utilizing Zonation 5 software. Two complementary prioritization algorithms were applied: Core Area Zonation (CAZ2), which emphasizes high-value biodiversity hotspots (regional complementarity), and the Additive Benefit Function (ABF), which prioritizes balanced representation across the landscape (local representativeness). Panels are organized by algorithm and metric: (a–f) PCAs based on Taxonomic Diversity (TD) using CAZ2. (g–l) PCAs based on Phylogenetic Diversity (PD) using CAZ2. (m–r) PCAs based on TD using ABF. (s–x) PCAs based on PD using ABF. Within each group, panels correspond to: all vascular plants, climbers, epiphytes, herbs, shrubs, and trees, respectively. Colored areas indicate the top 30% priority locations selected by the respective models.

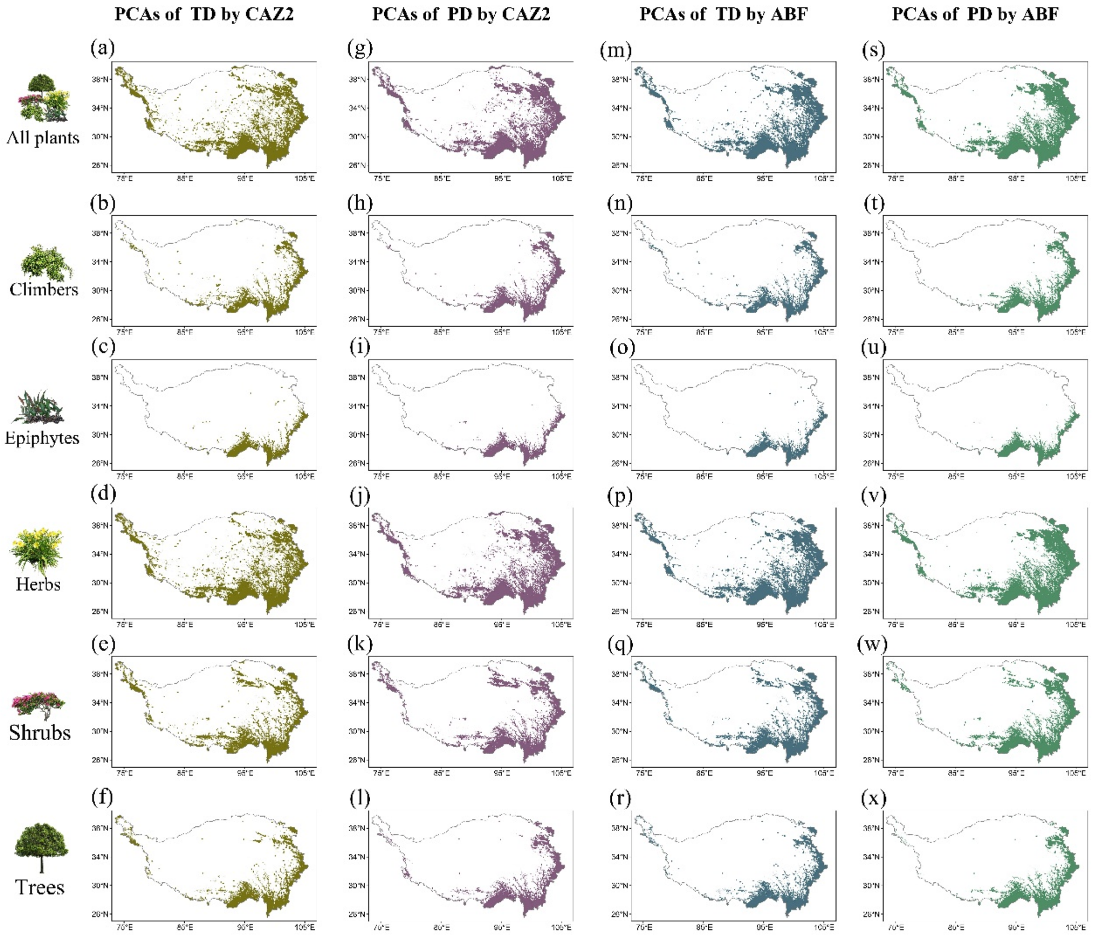


**Fig. S8. Spatial patterns of priority conservation areas (PCAs) across plant growth forms under an unweighted prioritization scheme.** This figure presents the sensitivity analysis corresponding to Supplementary Fig. 7, utilizing an unweighted prioritization scheme in Zonation 5 (i.e., uniform weight assigned to all species). As in the weighted analysis, PCAs were identified under a 30% area-protection target using two algorithms: CAZ2 (a–l) and ABF (m–x). Panels are organized by diversity metric and algorithm: (a–f) Taxonomic Diversity (TD) by CAZ2. (g–l) Phylogenetic Diversity (PD) by CAZ2. (m–r) TD by ABF. (s–x) PD by ABF. Within each set, panels follow the sequence: all vascular plants, climbers, epiphytes, herbs, shrubs, and trees. Note the differences in spatial configurations compared to the weighted scenarios in Supplementary Fig. 7, highlighting the influence of range-size rarity on conservation priorities.

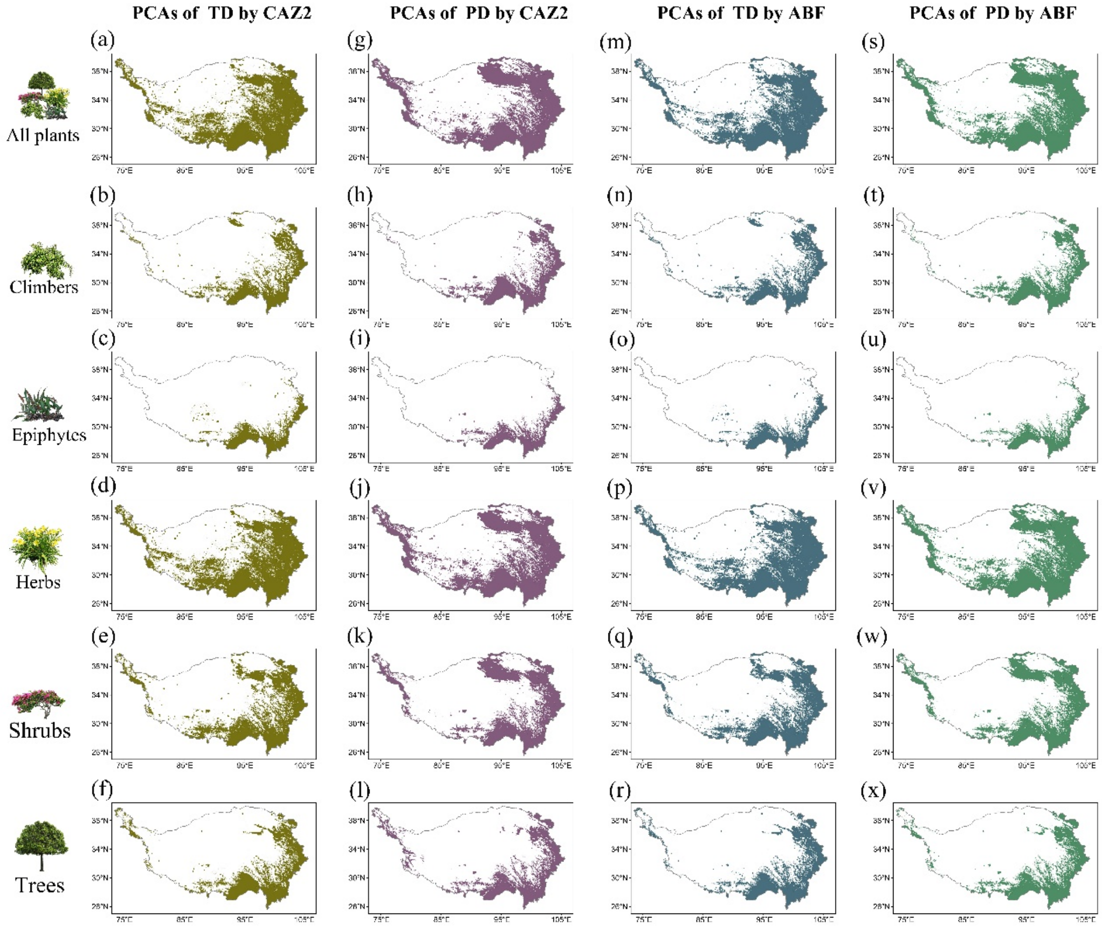


**Fig. S9. Spatial patterns of priority conservation areas (PCAs) under an expanded 50% area-protection target.** This figure illustrates conservation priorities identified under a 50% area-protection target (consistent with "Half-Earth" scenarios), using the weighted prioritization scheme in Zonation 5. The methodology mirrors Supplementary Fig. 7, applying two complementary algorithms: CAZ2 (Regional Complementarity) and ABF (Local Representativeness). Panels are grouped by algorithm and metric: (a–f) Taxonomic Diversity (TD) by CAZ2. (g–l) Phylogenetic Diversity (PD) by CAZ2. (m–r) TD by ABF. (s–x) PD by ABF. Within each group, panels follow the sequence: all vascular plants, climbers, epiphytes, herbs, shrubs, and trees. Comparing these patterns with Supplementary Fig. 7 (30% target) reveals the stability of core conservation areas as the protection network expands.

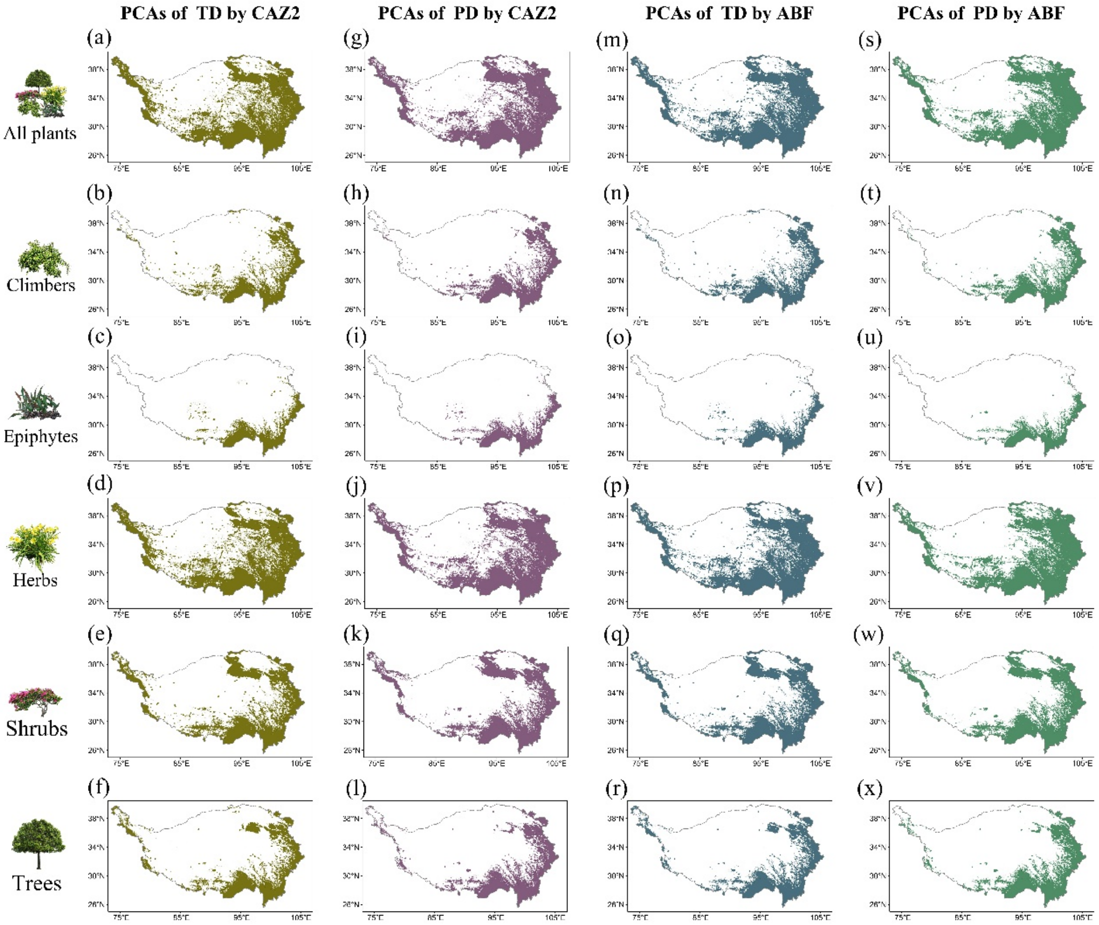


**Fig. S10. Spatial patterns of priority conservation areas (PCAs) under a 50% area-protection target with unweighted prioritization.** This figure presents the final sensitivity analysis, combining an expanded 50% area-protection target with an unweighted prioritization scheme (uniform species weights) in Zonation 5. The layout mirrors previous figures, facilitating comparison with the weighted 50% scenario (Supplementary Fig. 9) and the unweighted 30% scenario (Supplementary Fig. 8). Panels are organized by prioritization algorithm and diversity metric: (a–f) Taxonomic Diversity (TD) by CAZ2. (g–l) Phylogenetic Diversity (PD) by CAZ2. (m–r) TD by ABF. (s–x) PD by ABF. Within each group, panels correspond to the sequence: all vascular plants, climbers, epiphytes, herbs, shrubs, and trees.

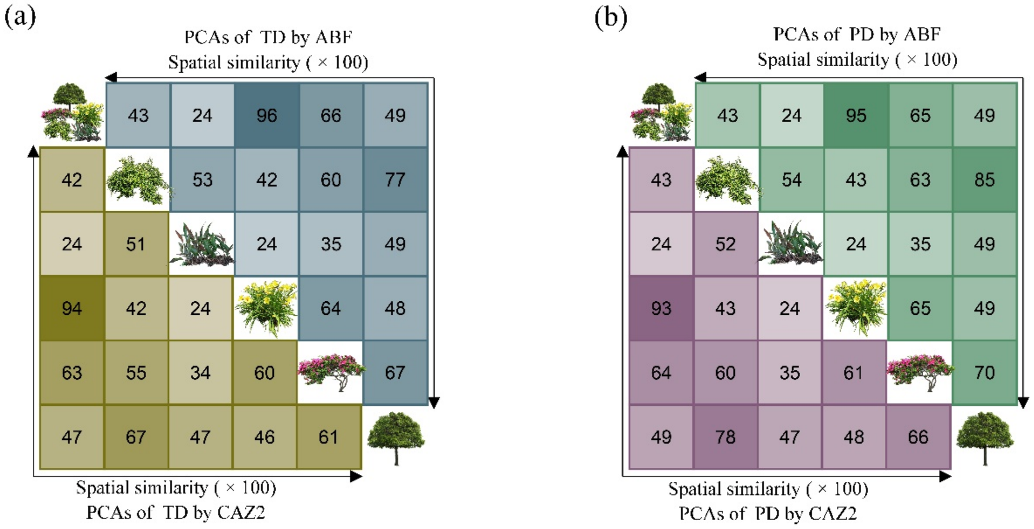


**Fig. S11. Pairwise spatial similarity of priority conservation areas (PCAs) across plant growth forms under a 30% unweighted target on the Qinghai-Tibet Plateau.** PCAs were identified using an unweighted prioritization scheme in Zonation 5 with a 30% area-protection target. Spatial overlap between PCA sets was quantified using the Jaccard similarity index. (a) Spatial similarity matrices for Taxonomic Diversity (TD). The lower triangle displays pairwise comparisons between growth forms identified by the CAZ2 algorithm (regional complementarity), while the upper triangle displays comparisons identified by the ABF algorithm (local representativeness). (b) Spatial similarity matrices for Phylogenetic Diversity (PD), following the same structure as (a): CAZ2 in the lower triangle and ABF in the upper triangle. In both panels, rows and columns follow the sequence: all vascular plants, climbers, epiphytes, herbs, shrubs, and trees. Darker colors indicate higher spatial similarity (higher Jaccard index) between the respective PCA sets.

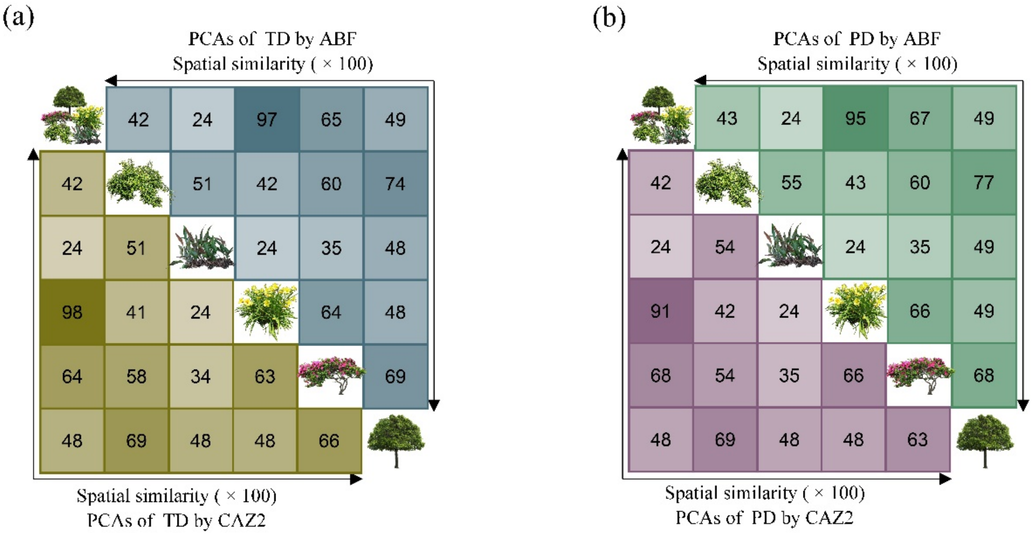


**Fig. S12. Pairwise spatial similarity of priority conservation areas (PCAs) across plant growth forms under a 50% weighted target on the Qinghai-Tibet Plateau.** PCAs were identified using an unweighted prioritization scheme in Zonation 5 with a 30% area-protection target. Spatial overlap between PCA sets was quantified using the Jaccard similarity index. (a) Spatial similarity matrices for Taxonomic Diversity (TD). The lower triangle displays pairwise comparisons between growth forms identified by the CAZ2 algorithm (regional complementarity), while the upper triangle displays comparisons identified by the ABF algorithm (local representativeness). (b) Spatial similarity matrices for Phylogenetic Diversity (PD), following the same structure as (a): CAZ2 in the lower triangle and ABF in the upper triangle. In both panels, rows and columns follow the sequence: all vascular plants, climbers, epiphytes, herbs, shrubs, and trees. Darker colors indicate higher spatial similarity (higher Jaccard index) between the respective PCA sets.

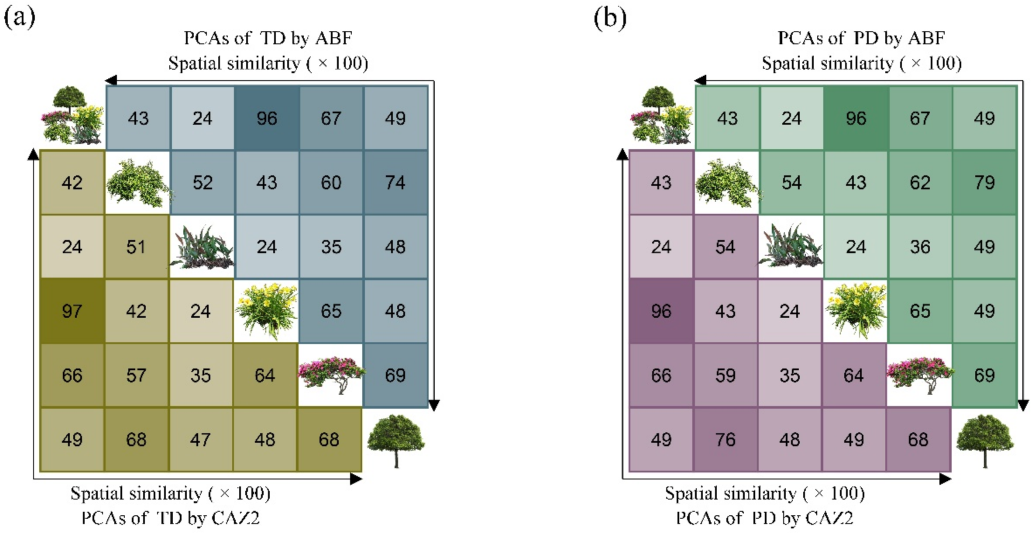


**Fig. S13. Pairwise spatial similarity of priority conservation areas (PCAs) across plant growth forms under a 50% unweighted target on the Qinghai-Tibet Plateau.** PCAs were identified using an unweighted prioritization scheme in Zonation 5 with a 30% area-protection target. Spatial overlap between PCA sets was quantified using the Jaccard similarity index. (a) Spatial similarity matrices for Taxonomic Diversity (TD). The lower triangle displays pairwise comparisons between growth forms identified by the CAZ2 algorithm (regional complementarity), while the upper triangle displays comparisons identified by the ABF algorithm (local representativeness). (b) Spatial similarity matrices for Phylogenetic Diversity (PD), following the same structure as (a): CAZ2 in the lower triangle and ABF in the upper triangle. In both panels, rows and columns follow the sequence: all vascular plants, climbers, epiphytes, herbs, shrubs, and trees. Darker colors indicate higher spatial similarity (higher Jaccard index) between the respective PCA sets.

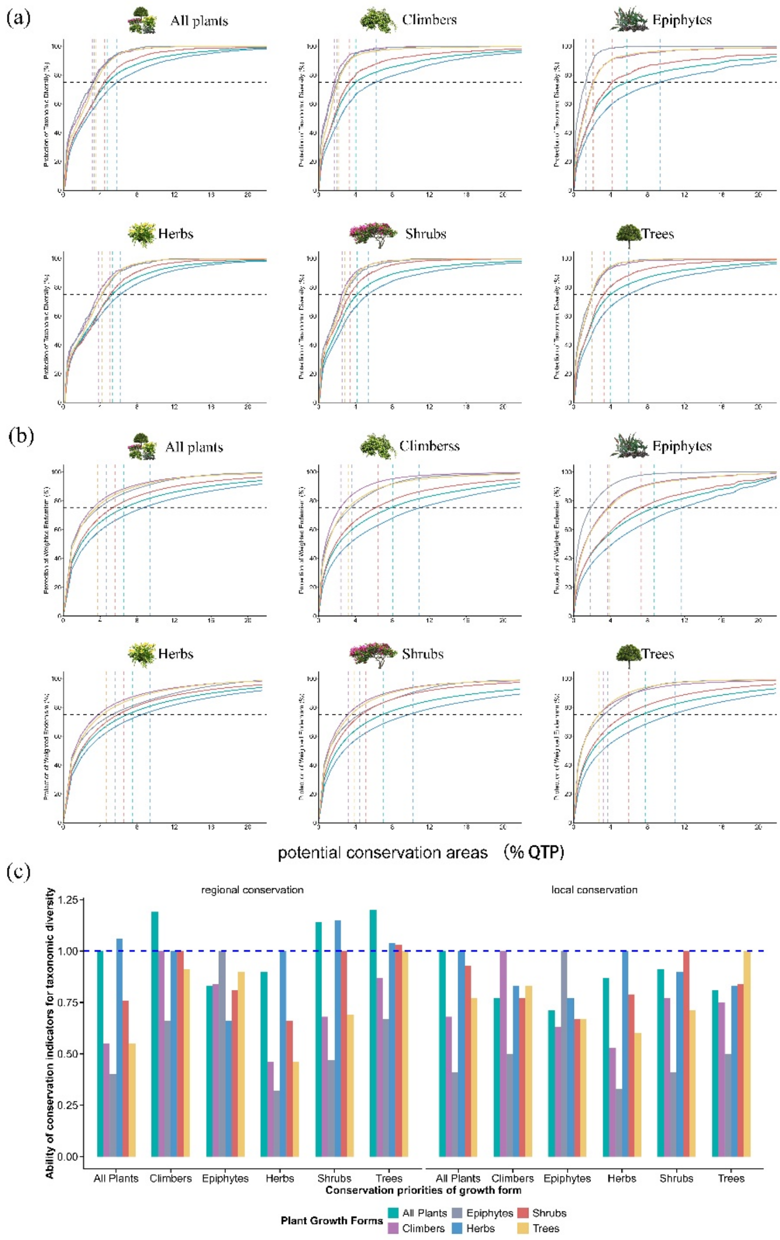


**Fig. S14. Effectiveness of growth forms as conservation surrogates for taxonomic diversity and weighted endemism.** We evaluated cross-taxon surrogacy, quantifying how efficiently conservation priorities identified for one growth form (surrogate) protect the diversity of others (targets). (a–b) Performance curves illustrating the accumulation of protected diversity as conservation areas expand based on surrogate priorities. Two conservation objectives were tested: (a) Regional Taxonomic Diversity (TD; proportion of total species richness protected) and (b) Local Weighted Endemism (WE; cumulative weighted endemism captured). Grey horizontal dashed lines indicate a 75% protection target, while vertical dashed lines denote the minimum land area required to achieve this target under each prioritization scheme. (c) Quantitative performance of each growth form as a surrogate, measured by the Ability of Conservation Indicator (ACI) for taxonomic diversity. The blue horizontal dashed line denotes ACI = 1 (indicating perfect surrogacy). All analyses are based on Zonation 5 using unweighted prioritization schemes. Color coding differentiates growth forms: climbers (purple), epiphytes (grey), herbs (blue), shrubs (red), trees (yellow), and all vascular plants (green).

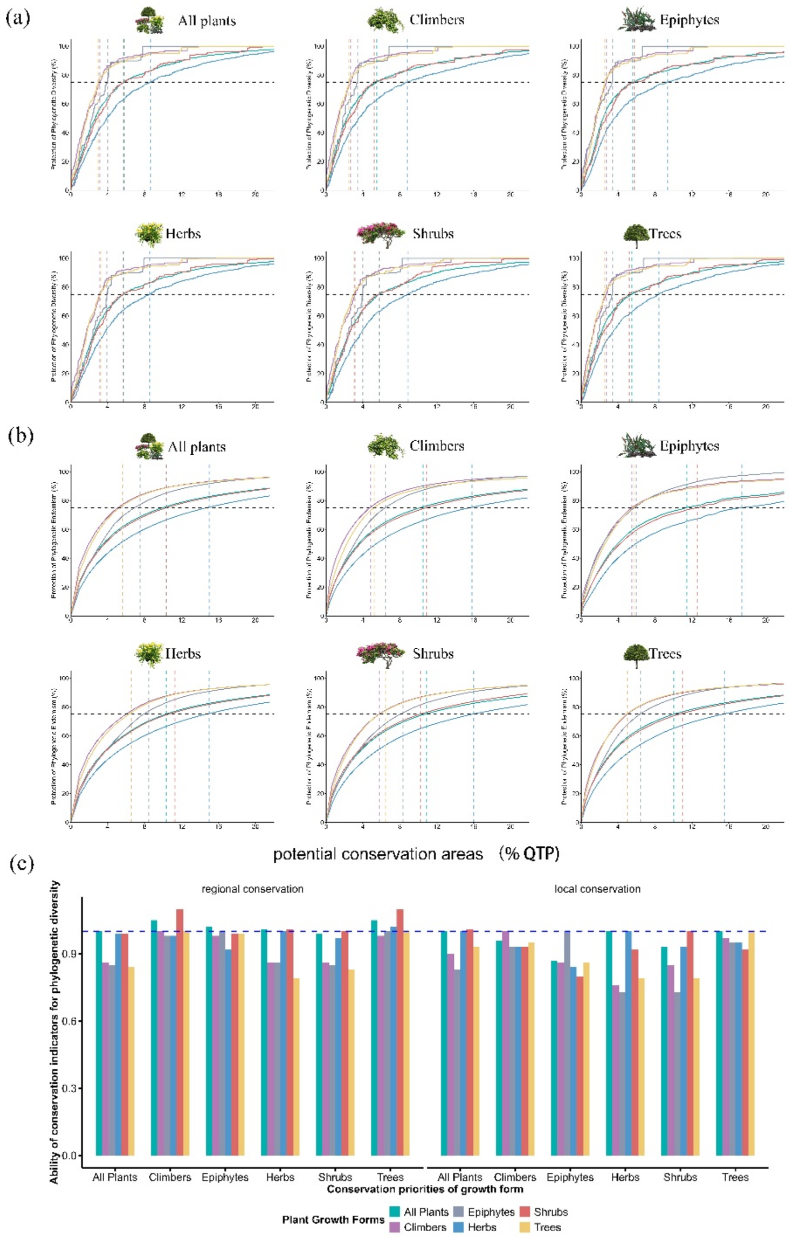


**Fig. S15. Effectiveness of growth forms as conservation surrogates for phylogenetic diversity and endemism.** This figure complements Supplementary Fig. 15 by evaluating cross-taxon surrogacy for phylogenetic dimensions using an unweighted prioritization scheme. (a–b) Performance curves showing the accumulation of protected diversity. Two conservation objectives were tested: (a) Regional Phylogenetic Diversity (PD; proportion of total evolutionary history protected) and (b) Local Phylogenetic Endemism (PE; retention of endemic branch lengths). Grey horizontal dashed lines indicate the 75% protection target; vertical dashed lines denote the minimum land area required to achieve it. (c) Quantitative performance of each growth form as a surrogate, measured by the Ability of Conservation Indicator (ACI) for taxonomic diversity. The blue horizontal dashed line denotes ACI = 1 (indicating perfect surrogacy). All analyses are based on Zonation 5 using weighted prioritization schemes. Color coding differentiates growth forms: climbers (purple), epiphytes (grey), herbs (blue), shrubs (red), trees (yellow), and all vascular plants (green).

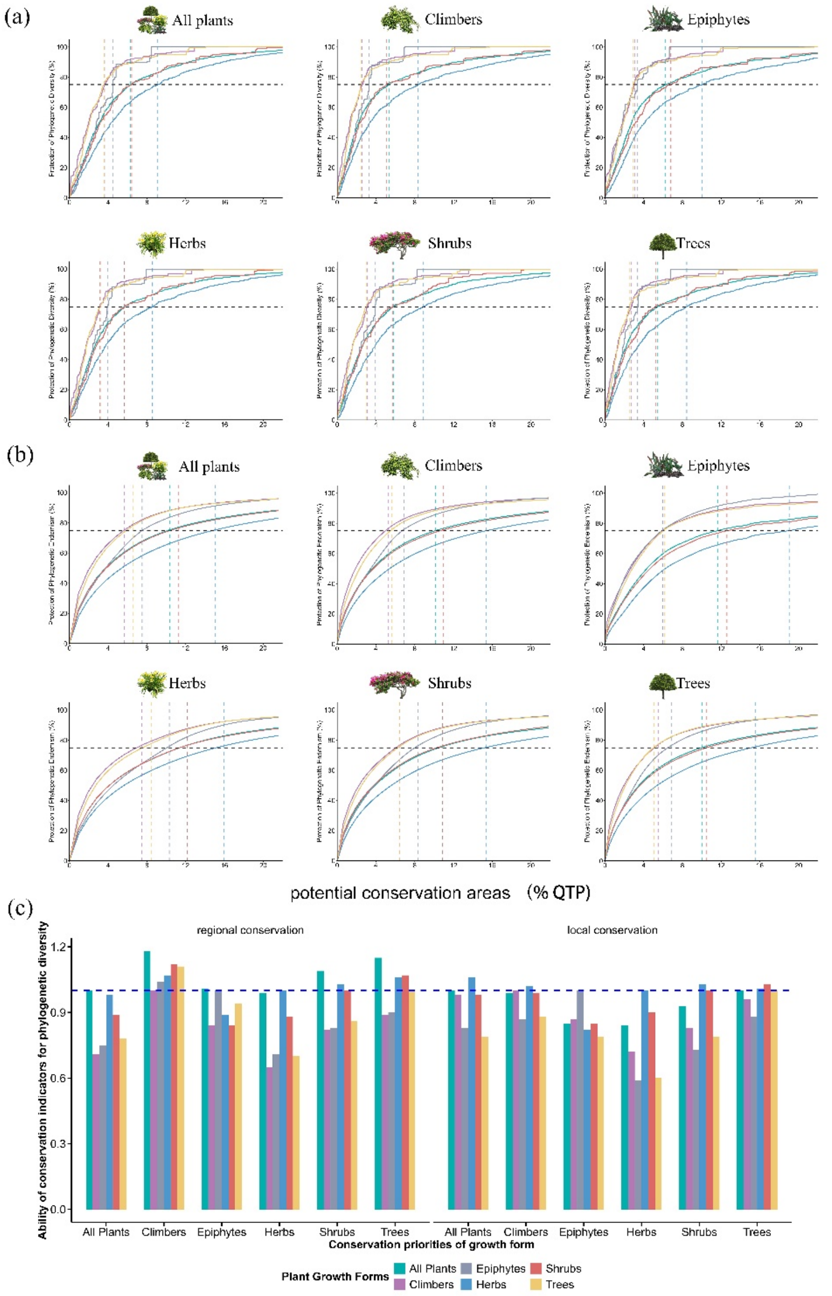


**Fig. S16. Effectiveness of growth forms as conservation surrogates for phylogenetic diversity under unweighted prioritization.** We evaluated how well the conservation priorities identified for one growth form (or all vascular plants) protect the diversity of all other growth forms. Two conservation objectives were tested: (a) representing regional phylogenetic diversity (PD; proportion of total phylogenetic diversity protected) and (b) capturing local phylogenetic endemism (PE; retention of endemic branch lengths). Curves show accumulation of protected diversity as simulated conservation areas expand according to each set of priorities. Grey horizontal dashed lines indicate the 75% protection target; vertical dashed lines show the minimum land area required to reach it. (c) Performance of each growth form as a surrogate for others was evaluated by quantifying the Ability of Conservation Indicator (ACI) for phylogenetic diversity. The blue horizontal dashed line denotes ACI = 1. All analyses are colour-coded by growth form: purple, climbers; grey, epiphytes; blue, herbs; red, shrubs; yellow, trees; green, all vascular plants. Conservation priorities were calculated using Zonation 5 with unweighted prioritization schemes.

**Table. S1. Taxonomic summary and occurrence data statistics for plant growth forms on the Qinghai-Tibet Plateau (QTP).** The table presents the number of families, genera, species, and total occurrence records for the entire flora and each specific growth form. Groups are arranged in descending order of data availability (occurrence frequency): herbs (most abundant), shrubs, trees, climbers, and epiphytes.

| Plant growth forms | Occurrence records count per species | | | | | | | | | | | |
|---|---|---|---|---|---|---|---|---|---|---|---|---|
| | >0 | | | | ≥5 | | | | 1~4 | | | |
| | family | genus | species | count | family | genus | species | count | family | genus | species | count |
| Herbs | 159 | 1298 | 8936 | 212402 | 145 | 1082 | 5966 | 206000 | 123 | 728 | 2970 | 6402 |
| Shrubs | 94 | 378 | 2088 | 48781 | 82 | 296 | 1436 | 47341 | 74 | 223 | 652 | 1440 |
| Trees | 118 | 431 | 1776 | 36294 | 105 | 332 | 1225 | 35058 | 89 | 249 | 551 | 1236 |
| Climbers | 75 | 271 | 1061 | 20141 | 63 | 205 | 669 | 19240 | 61 | 171 | 392 | 901 |
| Epiphytes | 32 | 120 | 607 | 7598 | 29 | 83 | 341 | 7053 | 19 | 83 | 266 | 545 |
| All Plants | 264 | 2150 | 14468 | 325216 | 243 | 1745 | 9637 | 314692 | 213 | 1281 | 4831 | 10524 |

**Table. S2. Summary of species distribution model (SDM) performance metrics for plant growth forms on the Qinghai-Tibet Plateau.** The table presents the validation statistics for the ensemble models constructed. Performance was evaluated using three complementary metrics: the Area Under the Receiver Operating Characteristic Curve (AUC), the True Skill Statistic (TSS), and the Continuous Boyce Index (Boyce). Values represent the mean ± standard deviation across all modeled species within each group. Species occupying fewer than five grid cells were excluded from the modeling process to ensure robust parameter estimation.

| Plant growth forms | AUC | TSS | Boyce |
|---|---|---|---|
| Climbers | 0.99±0.02 | 0.95±0.05 | 0.74±0.23 |
| Epiphytes | 0.99±0.01 | 0.95±0.05 | 0.72±0.25 |
| Herbs | 0.98±0.04 | 0.91±0.09 | 0.71±0.25 |
| Shrubs | 0.99±0.02 | 0.93±0.06 | 0.73±0.23 |
| Trees | 0.99±0.02 | 0.95±0.05 | 0.72±0.23 |
| All plants | 0.98±0.03 | 0.92±0.06 | 0.71±0.25 |

**Table. S3. Statistical breakdown of species mapped using Species Distribution Models (SDMs) versus circular buffers (Hybrid Mapping Strategy).** Two complementary approaches were employed to generate distribution ranges. (1) SDM-based ranges: Retained for species with robust model performance, defined as simultaneously meeting three criteria: AUC $\geq$ 0.7, TSS $\geq$ 0.5, and Boyce Index $\geq$ 0.4. (2) Buffer-based ranges: Applied to species with insufficient occurrence data (< 5 records) or those failing to meet the aforementioned SDM performance thresholds. For this latter group, ranges were delineated using a 10-km circular buffer around each occurrence point.

| Plant growth forms | Effective SDMs | | | Ineffective SDMs | | | Species of 10km buffer range | | |
|---|---|---|---|---|---|---|---|---|---|
| | family | genus | species | family | genus | species | family | genus | species |
| Herbs | 141 | 1012 | 5344 | 82 | 314 | 622 | 131 | 821 | 3592 |
| Shrubs | 82 | 282 | 1302 | 42 | 78 | 134 | 79 | 249 | 786 |
| Trees | 102 | 302 | 1097 | 43 | 84 | 128 | 93 | 283 | 679 |
| Climbers | 62 | 196 | 613 | 24 | 41 | 56 | 62 | 182 | 448 |
| Epiphytes | 28 | 78 | 300 | 13 | 29 | 41 | 22 | 92 | 307 |
| All Plants | 237 | 1637 | 8656 | 119 | 426 | 981 | 224 | 1426 | 5835 |

**Table. S4. Relationship between multi-faceted plant diversity and latitude/elevation on the Qinghai-Tibet Plateau modelled with trend surfaces.** Analysis was performed using ANOVA (DF: degrees of freedom; F: variance ratio; P: p-value; adjusted $R^2$: effect size). Diversity facets include species richness (SR), weighted endemism (WE), phylogenetic diversity (PD), and phylogenetic endemism (PE).

| Plant growth forms | Relationships of each diversity aspect with absolute latitude and elevation | DF | F | P | adjusted $R^2$ |
|---|---|---|---|---|---|
| All plants | SR ~ Latitude | 3534958 | 379857.9 | $<2 \times 10^{-16}$ | 0.23 |
| All plants | SR ~ Elevation | 3534958 | 748097.4 | $<2 \times 10^{-16}$ | 0.37 |
| Climbers | SR ~ Latitude | 1522938 | 343428.7 | $<2 \times 10^{-16}$ | 0.21 |
| Climbers | SR ~ Elevation | 1522938 | 749622.5 | $<2 \times 10^{-16}$ | 0.37 |
| Epiphytes | SR ~ Latitude | 862327 | 762866.2 | $<2 \times 10^{-16}$ | 0.38 |
| Epiphytes | SR ~ Elevation | 862327 | 703876.8 | $<2 \times 10^{-16}$ | 0.36 |
| Herbs | SR ~ Latitude | 3529429 | 321050.7 | $<2 \times 10^{-16}$ | 0.2 |
| Herbs | SR ~ Elevation | 3529429 | 659130 | $<2 \times 10^{-16}$ | 0.34 |
| Shrubs | SR ~ Latitude | 2413999 | 386957.6 | $<2 \times 10^{-16}$ | 0.24 |
| Shrubs | SR ~ Elevation | 2413999 | 686951.5 | $<2 \times 10^{-16}$ | 0.35 |
| Trees | SR ~ Latitude | 1721168 | 352327.7 | $<2 \times 10^{-16}$ | 0.22 |
| Trees | SR ~ Elevation | 1721168 | 692589.2 | $<2 \times 10^{-16}$ | 0.36 |
| All plants | PD ~ Latitude | 3534958 | 379715.3 | $<2 \times 10^{-16}$ | 0.23 |
| All plants | PD ~ Elevation | 3534958 | 1165641 | $<2 \times 10^{-16}$ | 0.48 |
| Climbers | PD~ Latitude | 1522938 | 514721.7 | $<2 \times 10^{-16}$ | 0.29 |
| Climbers | PD ~ Elevation | 1522938 | 999584.4 | $<2 \times 10^{-16}$ | 0.44 |
| Epiphytes | PD ~ Latitude | 862327 | 966975.7 | $<2 \times 10^{-16}$ | 0.44 |
| Epiphytes | PD ~ Elevation | 862327 | 843388.8 | $<2 \times 10^{-16}$ | 0.4 |
| Herbs | PD ~ Latitude | 3529429 | 552515.1 | $<2 \times 10^{-16}$ | 0.31 |
| Herbs | PD ~ Elevation | 3529429 | 1063725 | $<2 \times 10^{-16}$ | 0.46 |
| Shrubs | PD ~ Latitude | 2413999 | 490555.1 | $<2 \times 10^{-16}$ | 0.28 |
| Shrubs | PD ~ Elevation | 2413999 | 935198.5 | $<2 \times 10^{-16}$ | 0.43 |
| Trees | PD ~ Latitude | 1721168 | 494105.7 | $<2 \times 10^{-16}$ | 0.28 |
| Trees | PD ~ Elevation | 1721168 | 932573.2 | $<2 \times 10^{-16}$ | 0.43 |
| All plants | WE ~ Latitude | 3534958 | 136232.4 | $<2 \times 10^{-16}$ | 0.1 |
| All plants | WE ~ Elevation | 3534958 | 152617.6 | $<2 \times 10^{-16}$ | 0.11 |
| Climbers | WE ~ Latitude | 1522938 | 73353.55 | $<2 \times 10^{-16}$ | 0.06 |
| Climbers | WE ~ Elevation | 1522938 | 67385.74 | $<2 \times 10^{-16}$ | 0.05 |
| Epiphytes | WE ~ Latitude | 862327 | 131595.5 | $<2 \times 10^{-16}$ | 0.09 |
| Epiphytes | WE ~ Elevation | 862327 | 138287.7 | $<2 \times 10^{-16}$ | 0.1 |
| Herbs | WE ~ Latitude | 3529429 | 87844.97 | $<2 \times 10^{-16}$ | 0.07 |

| | | | | | |
|---|---|---|---|---|---|
| Herbs | WE ~ Elevation | 3529429 | 127246.2 | $<2 \times 10^{-16}$ | 0.09 |
| Shrubs | WE ~ Latitude | 2413999 | 99058.56 | $<2 \times 10^{-16}$ | 0.07 |
| Shrubs | WE ~ Elevation | 2413999 | 113846.5 | $<2 \times 10^{-16}$ | 0.08 |
| Trees | WE ~ Latitude | 1721168 | 139263.2 | $<2 \times 10^{-16}$ | 0.1 |
| Trees | WE ~ Elevation | 1721168 | 170219.7 | $<2 \times 10^{-16}$ | 0.12 |
| All plants | PE ~ Latitude | 3534958 | 139263.2 | $<2 \times 10^{-16}$ | 0.1 |
| All plants | PE ~ Elevation | 3534958 | 170219.7 | $<2 \times 10^{-16}$ | 0.12 |
| Climbers | PE ~ Latitude | 1522938 | 61197.37 | $<2 \times 10^{-16}$ | 0.05 |
| Climbers | PE ~ Elevation | 1522938 | 55012.43 | $<2 \times 10^{-16}$ | 0.04 |
| Epiphytes | PE ~ Latitude | 862327 | 88051.23 | $<2 \times 10^{-16}$ | 0.07 |
| Epiphytes | PE ~ Elevation | 862327 | 127684.7 | $<2 \times 10^{-16}$ | 0.09 |
| Herbs | PE ~ Latitude | 3529429 | 131595.5 | $<2 \times 10^{-16}$ | 0.09 |
| Herbs | PE ~ Elevation | 3529429 | 138287.7 | $<2 \times 10^{-16}$ | 0.1 |
| Shrubs | PE ~ Latitude | 2413999 | 99058.55 | $<2 \times 10^{-16}$ | 0.07 |
| Shrubs | PE ~ Elevation | 2413999 | 113846.5 | $<2 \times 10^{-16}$ | 0.08 |
| Trees | PE ~ Latitude | 1721168 | 136311.2 | $<2 \times 10^{-16}$ | 0.1 |
| Trees | PE ~ Elevation | 1721168 | 152728 | $<2 \times 10^{-16}$ | 0.11 |

**Table. S5. Relationship between taxonomic and phylogenetic composition and diversity of plant assemblages across growth forms on the Qinghai-Tibet Plateau.** Analyses were conducted for all vascular plants and five growth forms within 1-km² grid cells. Associations between composition and diversity metrics—species richness (SR) and phylogenetic diversity (PD)—were evaluated using Spearman's rank correlation (ρ).

| | | Species richness | | | | | | Phylogenetic diversity | | | | | |
|---|---|---|---|---|---|---|---|---|---|---|---|---|---|
| | | All plants | Climbers | Epiphytes | Herbs | Shrubs | Trees | All plants | Climbers | Epiphytes | Herbs | Shrubs | Trees |
| Taxonomic Composition | Climbers | 0.77 | 0.98 | 0.69 | 0.76 | 0.72 | 0.74 | 0.77 | 0.98 | 0.69 | 0.77 | 0.71 | 0.74 |
| | Epiphytes | 0.66 | 0.71 | 0.99 | 0.65 | 0.68 | 0.74 | 0.69 | 0.71 | 0.99 | 0.68 | 0.67 | 0.74 |
| | Herbs | -0.75 | -0.69 | -0.65 | -0.71 | -0.91 | -0.82 | -0.78 | -0.69 | -0.65 | -0.74 | -0.91 | -0.82 |
| | Shrubs | 0.67 | 0.58 | 0.57 | 0.64 | 0.89 | 0.72 | 0.71 | 0.57 | 0.57 | 0.66 | 0.89 | 0.72 |
| | Trees | 0.75 | 0.7 | 0.69 | 0.72 | 0.8 | 0.95 | 0.79 | 0.69 | 0.69 | 0.75 | 0.8 | 0.94 |
| Phylogenetic Composition | Climbers | 0.8 | 0.99 | 0.71 | 0.8 | 0.75 | 0.78 | 0.8 | 0.99 | 0.71 | 0.8 | 0.74 | 0.78 |
| | Epiphytes | 0.65 | 0.69 | 0.99 | 0.64 | 0.67 | 0.73 | 0.68 | 0.69 | 0.99 | 0.67 | 0.66 | 0.72 |
| | Herbs | -0.75 | -0.67 | -0.62 | -0.72 | -0.87 | -0.83 | -0.79 | -0.67 | -0.62 | -0.74 | -0.88 | -0.83 |
| | Shrubs | 0.73 | 0.57 | 0.47 | 0.71 | 0.9 | 0.72 | 0.76 | 0.56 | 0.47 | 0.71 | 0.92 | 0.73 |
| | Trees | 0.81 | 0.75 | 0.71 | 0.8 | 0.85 | 0.97 | 0.85 | 0.74 | 0.71 | 0.81 | 0.84 | 0.98 |
